# Unlocking the soundscape of coral reefs with artificial intelligence: pretrained networks and unsupervised learning win out

**DOI:** 10.1101/2024.02.02.578582

**Authors:** Ben Williams, Santiago M. Balvanera, Sarab S. Sethi, Timothy A.C. Lamont, Jamaluddin Jompa, Mochyudho Prasetya, Laura Richardson, Lucille Chapuis, Emma Weschke, Andrew Hoey, Ricardo Beldade, Suzanne C. Mills, Anne Haguenauer, Frederic Zuberer, Stephen D. Simpson, David Curnick, Kate E. Jones

## Abstract

Passive acoustic monitoring can offer insights into the state of coral reef ecosystems at low-costs and over extended temporal periods. Comparison of whole soundscape properties can rapidly deliver broad insights from acoustic data, in contrast to the more detailed but time-consuming analysis of individual bioacoustic signals. However, a lack of effective automated analysis for whole soundscape data has impeded progress in this field. Here, we show that machine learning (ML) can be used to unlock greater insights from reef soundscapes. We showcase this on a diverse set of tasks using three biogeographically independent datasets, each containing fish community, coral cover or depth zone classes. We show supervised learning can be used to train models that can identify ecological classes and individual sites from whole soundscapes. However, we report unsupervised clustering achieves this whilst providing a more detailed understanding of ecological and site groupings within soundscape data. We also compare three different approaches for extracting feature embeddings from soundscape recordings for input into ML algorithms: acoustic indices commonly used by soundscape ecologists, a pretrained convolutional neural network (P-CNN) trained on 5.2m hrs of YouTube audio and a CNN trained on individual datasets (T-CNN). Although the T-CNN performs marginally better across the datasets, we reveal that the P-CNN is a powerful tool for identifying marine soundscape ecologists due to its strong performance, low computational cost and significantly improved performance over acoustic indices. Our findings have implications for soundscape ecology in any habitat.

**Author Summary:** Artificial intelligence has the potential to revolutionise bioacoustic monitoring of coral reefs. So far, a limited set of work has used machine learning to train detectors for specific sounds such as individual fish species. However, building detectors is a time-consuming process that involves manually annotating large amounts of audio followed by complicated model training, this must then be repeated all over again for any new dataset. Instead, we explore machine learning techniques for whole soundscape analysis, which compares the acoustic properties of raw recordings from the entire habitat. We identify multiple machine learning methods for whole soundscape analysis and rigorously test these using datasets from Indonesia, Australia and French Polynesia. Our key findings show use of a neural network pretrained on 5.2m hours of unrelated YouTube audio offers a powerful tool to produce compressed representations of reef audio data, conserving the data’s key properties whilst being executable on a standard personal laptop. These representations can then be used to explore patterns in reef soundscapes using “unsupervised machine learning”, which is effective at grouping similar recordings periods together and dissimilar periods apart. We show these groupings hold relationships with ground truth ecological data, including coral coverage, the fish community and depth.

## 1. Introduction

Effective monitoring of coral reefs is essential for supporting their conservation and restoration (1). Monitoring data are typically collected using diver-led underwater visual census surveys. However, these diver-led surveys incur high expertise, logistical and financial costs (1,2). Passive acoustic monitoring (PAM) presents an alternative means of gathering monitoring data which can be collected with greater ease and over extended periods (3,4). However, in contrast to ecosystems where PAM is well established, the species identity for any given sound in reef PAM recordings is usually unknown (5,6). Instead, coral reef PAM typically attempts to find relationships between the ecological community and its ‘soundscape’, the full extent of environmental sounds present (7). Information held within reef soundscapes often correlates with ecological community metrics (e.g., fish diversity, coral cover)(8–10) and better captures temporal trends or the presence of cryptic organisms than traditional methods (11–14).

Given the ease of PAM data collection, automated analysis is required to maximize the potential of these data. Previous attempts at automated analysis of whole soundscape data have primarily used acoustic indices, which are formulas designed to quantify the properties of a spectrogram relevant to natural soundscapes (e.g., total amplitude and entropy across time). However, comparisons of individual handcrafted features, such as acoustic indices provides a limited resolution (15,16). This is further compounded by a lack of standardisation for the many parameters required when calculating indices (16,17). Machine learning (ML) presents a powerful alternative for automating the analysis of bioacoustic data across multiple ecosystems and taxa, so far primarily focused on generating species detectors (18–20). However, applications of ML for soundscape ecology, rather than individual bioacoustic signals, are much less developed for most ecosystems and are almost entirely untested on coral reefs (21–23). The optimum ML approaches to employ and insights these approaches can generate remain unknown.

Two common types of ML are available to soundscape ecologists: supervised and unsupervised learning. Supervised learning involves training ML algorithms on labelled recordings, where labels likely correspond to an ecological metric or habitat category of interest. The trained algorithm can then be used to recognize and predict these categories in new, unseen data. An alternative is unsupervised learning, which does not require labelled data. Instead, unsupervised algorithms can be used to reveal patterns and structures within the data by grouping similar recordings using clustering algorithms. The outputs from unsupervised learning can be used in a semi-supervised way to label new recordings that are grouped closely with labelled recordings, used to reveal temporal patterns within sites, find anomalous periods, and more (24).

Both supervised and unsupervised ML algorithms leverage feature embeddings, compressed representations of the data either calculated using handcrafted features designed by experts or learned by the algorithm during the process. We identified three approaches broadly representative of the techniques that can be used to extract feature embeddings from soundscape data, which remain largely unbenchmarked against one another (25). The first approach uses a ‘compound index’, generated by calculating multiple acoustic indices from recordings and combining these indices into a vector. A limitation of handcrafted features is that they often overlook other valuable but unidentified properties. Our second approach therefore utilised deep learning models. Deep learning uses neural networks to learn the most relevant features from complex data types such as audio (15). A disadvantage of deep learning is that greater computational resources and expertise are required to train these networks (26). Our third approach attempted to retain the benefits of deep learning while overcoming its disadvantages by using a pretrained neural network. Pretrained networks are typically trained on large datasets to predict a broad selection of classes. If the final classification layers are removed, these networks can be used as computationally inexpensive embedding extractors that generalize well to new data; this process is known as ‘transfer learning’. Like a compound index, these embeddings can be used to train lower-complexity ML algorithms, making this process significantly faster than training a custom neural network from scratch (27).

In this study, we tested the ability of supervised and unsupervised ML algorithms to identify meaningful ecological groupings from coral reef soundscape recordings (Fig. 1a). Additionally, we compared the suitability of the three feature embedding extraction approaches we identified: a compound index, an industry standard pretrained neural network (P-CNN) trained on 5.2m hrs of unrelated YouTube audio (VGGish)(28), and the same network custom trained on reef soundscape data (T-CNN)(Fig. 1a). Finally, we benchmarked our ML approaches against the predominant method for automated reef soundscape analysis of recent years: individual acoustic indices (17,29,30). To ensure that our results are representative of reef soundscapes broadly, we used recordings from three biogeographically distinct locations: Indonesia, Australia and French Polynesia (Fig. 1b)(31).

**Fig. 1.**
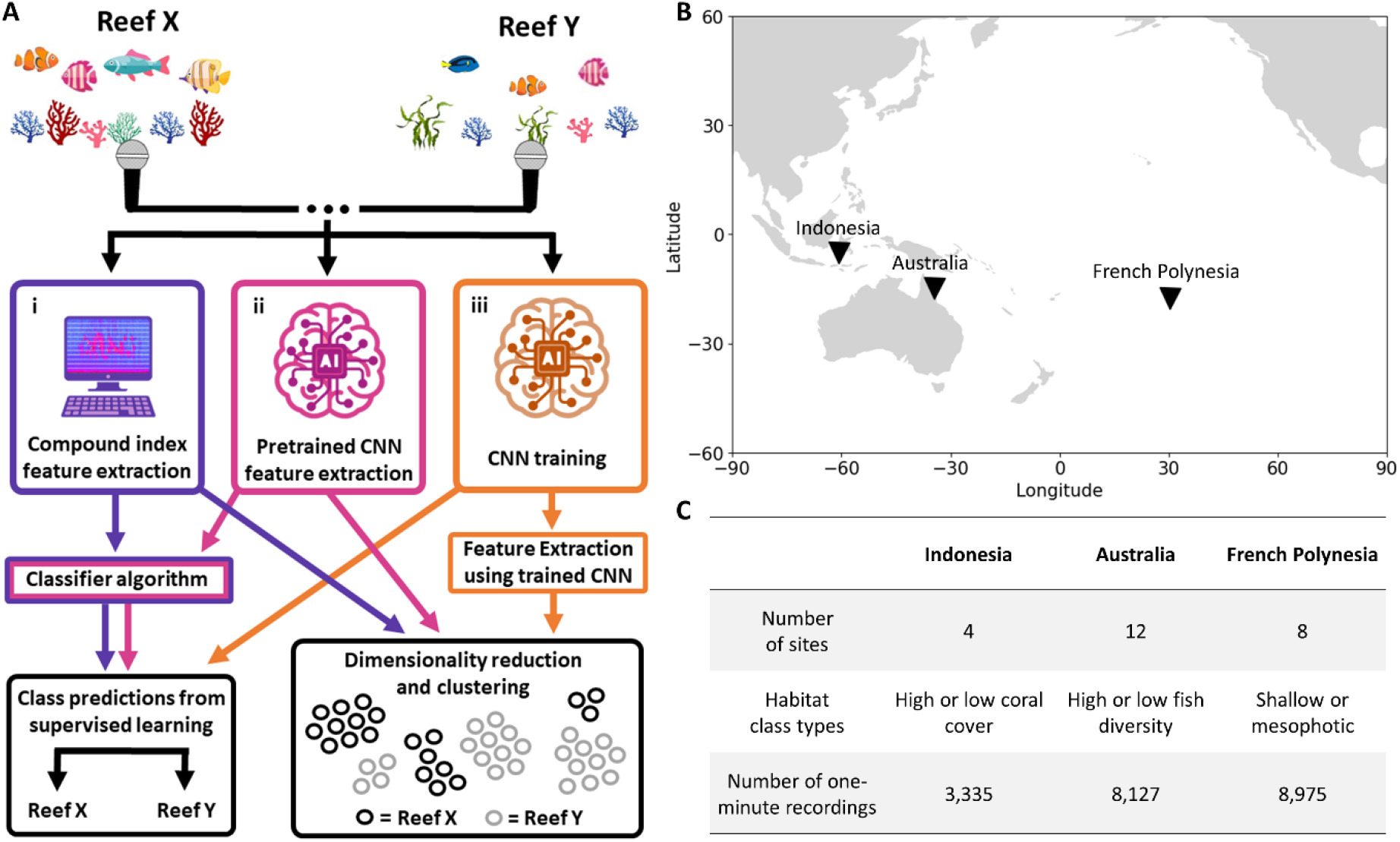
**(A)** Analytical workflow used on each coral reef soundscape dataset. The data from the reef soundscape datasets are entered into three pathways. For both the (i) compound index and (ii) pretrained CNN pathways this first involves a feature extraction stage. For supervised learning tasks these extracted feature embeddings are then entered into a classifier algorithm (random forest) trained to predict the class of new recordings. For unsupervised learning, dimensionality reduction and visualisation are performed on extracted embeddings using uniform manifold approximation (UMAP) followed by a clustering algorithm (affinity propagation). The (iii) trained CNN pathway performs supervised classification tasks directly and, once trained, is adapted and used as a feature extractor on recordings. Dimensionality reduction and clustering is then performed on these learned feature embeddings. **(B)** Geographic location of the datasets used. **(C)** Table of the key information from each dataset. The Indonesian sites were divided into high (91.2-93.1%) and low (2.1-17.6%) coral cover groups. The Australian sites were divided into a class with high biomass (20.5-52.2kg) and species richness (45spp - 53spp) scores per transect, and, low biomass (5.2-9.8kg) and species richness (30spp – 35spp) scores per transect (fig. S6). The French Polynesian sites were divided into shallow (10-15m) and mesophotic (55-65m) depth classes.

Each dataset included two unique habitat types: high or low coral cover for Indonesia, high or low fish diversity for Australia, and shallow or mesophotic depths for French Polynesia (Fig. 1c). The three datasets were treated independently throughout to provide a suite of unique challenges.

## 2. Results

### 2.1. Exploring coral reef soundscapes with unsupervised learning

Each dataset was divided into one-minute recording periods and embeddings were extracted from these data using the three methods: the compound index, P-CNN and T-CNN. Given that embedding must be learned for the T-CNN, these were obtained by training a separate network for each dataset to identify the sites within the dataset (given that the site origin should always be available for a recording), followed by using the networks to extract embeddings from their respective datasets.

The first unsupervised technique tested was uniform manifold approximation (UMAP)(32). This technique can reduce the dimensions of data to produce two-dimensional plots that group similar recordings closer together in the embedding space. Qualitative inspection of UMAP visualisations from each individual dataset revealed that the known habitat classes were frequently key drivers behind the clusters that were formed, indicating the influence of these on the soundscape (Fig. 2b; Fig. S1). Of the three embedding methods, the recordings were most clearly separated into discrete clusters that conformed to habitat classes by the T-CNN, followed by the P-CNN (Fig. 2b; Fig. S1). The UMAP plots also revealed a unique soundscape ‘fingerprint’ for many sites, with the French Polynesian dataset exhibiting the clearest discrete clusters for each site (Fig. 2b).

**Fig. 2.**
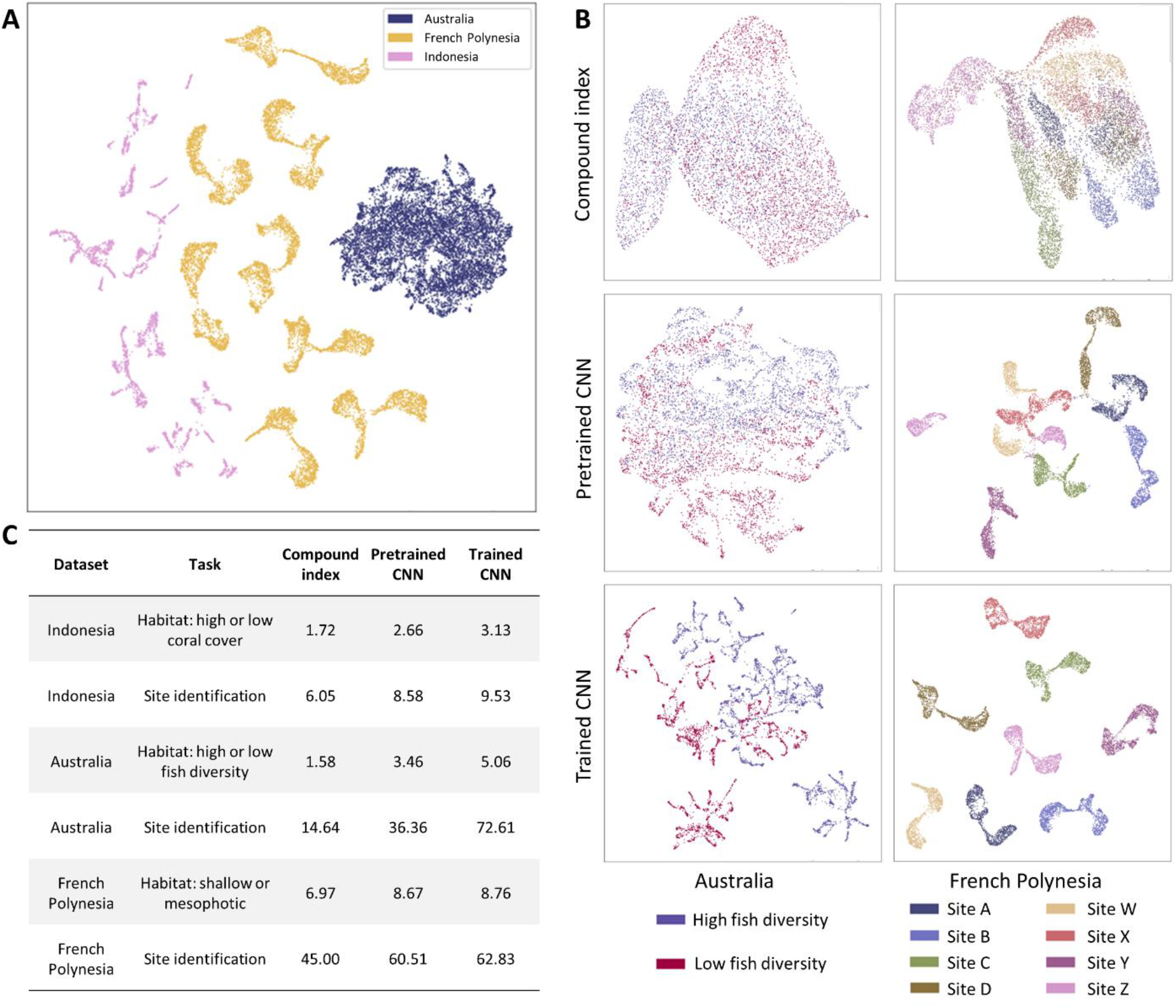
**(A)** Uniform manifold approximation (UMAP) dimensionality reduction and plotting applied to P-CNN embeddings of all 20,437 one-minute recordings (3,335 from Indonesia; 8,127 from Australia; 8,925 from French Polynesia). Individual points represent a one-minute recording. **(B)** Selected plots from uniform manifold approximation (UMAP) dimensionality reduction performed on the compound index, pretrained CNN and trained CNN embeddings. Plots were produced from the Australian and French Polynesian datasets and are labelled with colours corresponding to the habitat class (high or low fish diversity) and site, respectively. The remaining plots can be found in Supplementary 1. **(C)** Chi-squared scores (x 10^3^) generated from contingency tables of each recording’s true class and the cluster that each recording was assigned to by affinity propagation clustering. Affinity propagation was performed on UMAP embeddings generated using each of the three methods (compound index, pretrained CNN and trained CNNs). Higher scores indicate that clusters better represented true classes. All chi-square scores were highly significant (p<0.001).

In addition to identifying site and habitat classes, interactive UMAP plots also revealed temporal patterns within the data (Text S1; Supp. 2). The interactive plots showed that within each dataset, recordings from similar temporal periods typically grouped closer together. For example, crepuscular periods linked separate night and day clusters for the French Polynesian dataset, while the soundscape of two sites in Indonesia converged around the new moon (Text S1; Supp. 2). A visualisation of the three full datasets (Indonesia, Australia, French Polynesia) using the P-CNN UMAP embeddings was also performed. This plot showed that each formed distinct groups separate from the other datasets, indicating that the properties of the recordings from each location were unique from those of the other datasets (Fig. 2a).

Finally, a quantitative assessment of unsupervised performance was performed using affinity propagation clustering on UMAP embeddings. Chi-square tests were used to determine how well clusters split the data nonrandomly into ground-truth habitat and site classes. Across all three datasets, the T-CNNs once again reported the highest fidelity to both the habitat and site classes. In every instance, this was followed by the P-CNN and then compound index (Fig. 2c). However, the chi-square scores for the T-CNNs were only marginally better than those of the P-CNN for the French Polynesian habitat and site identification tasks (Fig. 2c). This was especially impressive for the site task given that the T-CNN embeddings were taken from networks trained to identify the sites, highlighting the strength of the P-CNN.

### 2.2. Predicting habitat class and site identity with supervised learning

For supervised learning, ML classifiers were trained on labelled recordings to directly predict the class of held-out test recordings, and the prediction accuracy on the test recordings is reported. To rigorously separate training and test data for the habitat classification tasks, we elected to exclude entire sites from training for the Australian and French Polynesian datasets, whereas only entire recording blocks (from instrument rotation) for the Indonesian set could be excluded due to low site replication (Text S2). For site classification, full recording blocks from each site (e.g., a contiguous 24 hr period) were held out for evaluation to mitigate against temporal autocorrelation. For each task, we extensively tested the influence of the train and test splits selected using either 32 or 100 alternative splits for each task (Text S2). Random forest classifiers were trained on the compound index or P-CNN embeddings, whereas class predictions were output directly from the T-CNNs.

Overall, the classifiers performed well at predicting habitat and individual site classes of one-minute reef soundscape recordings. Pooling classifications from these short windows over longer temporal periods therefore represents a potential route for soundscape ecologists to infer habitat states. The six tasks exhibited a range of difficulty, with a mean accuracy of 0.56 (±0.05) to 1.00 (±0.0) and performance above random classification ranging from 0.23 (±0.11) to 0.87 (±0.0)(Fig. 3; Table S1). The strongest performance at site classification tasks was reported for the French Polynesian dataset and the weakest was reported for the Australian dataset. However, we find the train and test divisions selected for a given repeat strongly influenced results. For example, the French Polynesian habitat classification task had a mean accuracy of 0.83 (±0.2) using the T-CNN, but some repeats failed to improve beyond random chance at discriminating classes.

**Fig. 3.**
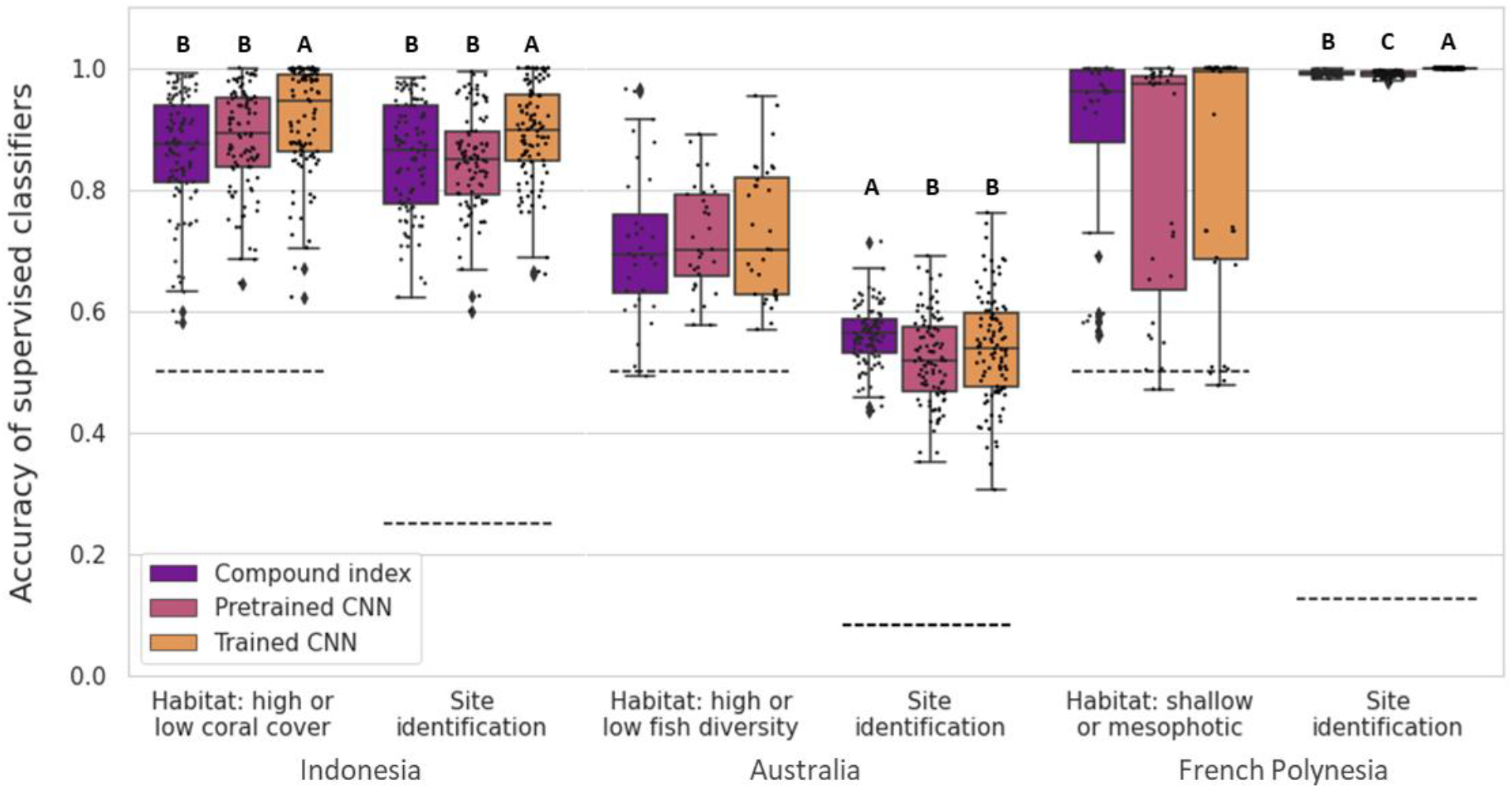
Boxplots of supervised classifier accuracies for the six different tasks across repeated training instances using each of the three embedding extraction methods (compound index, pretrained CNN and trained CNN). Boxes and their bars represent the 25^th^, 50^th^ and 75^th^ quartiles. The black dotted lines represent the expected accuracy using random classification (1 / number of classes). For each task, letters indicate a significant difference between these groupings according to ANOVA. The directions of significant differences reported by the Tukey HSD are indicated, with ‘A’ indicating the group with the highest accuracy, ‘B’ indicating the second highest accuracy, and where present, ‘C’ indicating the third group with the lowest accuracy. N = 100 for the number of repeats performed for all tasks, except for the ‘high or low fish diversity and ‘shallow or mesophotic’ tasks, where N = 32 (Text S2).

The three embedding approaches reported similar accuracies compared to one another for any given task. ANOVA tests reported no significant difference between any of the three methods for the habitat identification tasks set from the Australian and French Polynesian datasets (Fig. 3; Table S2). However, differences in accuracy were reported between each method for the Indonesian habitat classifier and for classifiers trained to identify sites for each dataset. For three of the four tasks where a difference was reported, the T-CNNs outperformed the other two methods: the habitat and site classifiers for the Indonesian dataset and the site identity classifier for the French Polynesian dataset. However, the compound index had significantly greater accuracy than the P-CNN and T-CNN for the Australian site classifier.

Inspection of the confusion matrices used to interpret classifier performance revealed that the three embedding extraction methods reported similar patterns of misclassification for each task, where samples from any given class were assigned to the same incorrect classes across methods (Fig. S2). The only exception was the French Polynesian habitat classification task, with 84.1 % and 67.6 % of the mesophotic recordings misclassified as shallow by the compound index and P-CNN respectively, while only 42.3 % with the T-CNN. The only instance where a class was misclassified the majority of the time across all repeats occurred for site A in the Australian dataset, where samples were more likely to be assigned to class G (Fig. S2). This misclassification was likely due to the close proximity of these sites (Fig. S5) and their habitat attributes, both of which were ‘low’ fish diversity sites.

### 2.3. Benchmarking against current automation: acoustic indices

Statistical comparisons of individual acoustic indices represent the predominant method for automated analysis of reef soundscape data indices (17,29,30). To assess the performance of individual indices on our data, we deliberately selected the individual acoustic indices that reported the most significant differences between each habitat class. Despite this biased selection, our analysis showed that these indices were still unable to classify individual recordings according to the accuracy of the ML (Fig 4). The proportions of recordings that could be unambiguously classified as the correct habitat type were only 5.2%, 13.0%, and 12.2% for each dataset respectively. Even the lowest-performing ML classifiers across all combinations of tasks and repeats achieved accuracies of 58.1%, 57.7% and 47.1% for each embedding type (Fig. 3). Furthermore, individual acoustic indices were unable to classify the site of origin for the recordings (Fig. S3).

**Fig. 4.**
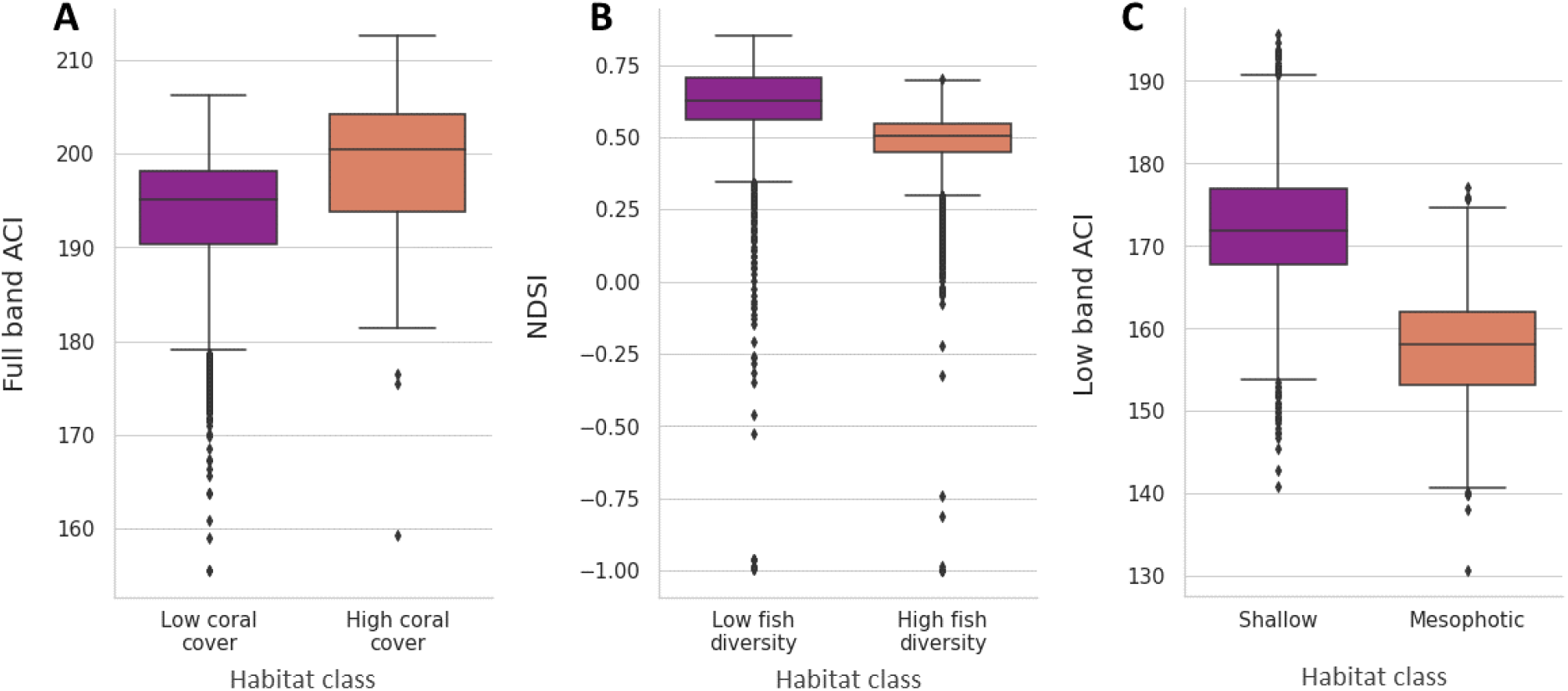
Boxplots of the acoustic index for which the highest significant difference between habitat classes was reported for the **(A)** Indonesian, **(B)** Australian and **(C)** French Polynesian datasets. These were the full-band acoustic complexity index (ACI), normalised difference soundscape index (NDSI), and low-band acoustic complexity index (ACI) respectively, which reported Mann-Whitney U test results of: u = 2.00 x 10^6^, p<0.0001, u = 7.71 x 10^6^, p<0.0001, and, u = 3.64 x 10^6^, p<0.0001. Boxes and their bars represent the 25th, 50th and 75th quartiles.

## 3. Discussion

### 3.1 Machine learning unlocks greater insights from coral reef soundscapes

Using three biogeographically independent datasets we demonstrate the potential of machine learning (ML) to automate the analysis of coral reef soundscape data at scale (20,437 one-minute recordings in the present study). Previous work using automated techniques has predominantly used individual acoustic indices, which have revealed relationships between the soundscape and key ecological attributes on reefs (10,33–36). However, comparing individual indices provides only a weak signal and often fails in new contexts (2,25,37,38); this finding is supported by our findings (Fig. 4, Fig. S4). Instead, we show that combining features into embeddings and inputting these into ML algorithms provides a powerful alternative, especially when using embeddings output by deep-learning models. However, care should be taken when designing the sampling regimes as well as dividing training and evaluation data.

Our results further support the hypothesis that the raw soundscapes of coral reefs share a relationship with ecological properties such as the fish community and coral cover. We also provide further evidence that shallow and mesophotic reefs often exhibit distinct soundscapes (39–41) (Fig. 3). Furthermore, ML was often able to differentiate individual sites, an insight which could not have been elucidated using acoustic indices (Fig. S3). However, our site identification findings underscore the need to mitigate recorder bias, which is frequently a factor in machine learning analyses. Notably, our French Polynesia dataset exhibited the most pronounced site identification trend, possibly because it was the only dataset in which the instruments could not be rotated (Text S2).

Our findings using unsupervised learning provide further evidence that temporal patterns influence reef soundscapes (5,13,42), with clusters typically grouping recordings from similar time periods (Text S1, Supp. 2). When considering the performance across datasets, the unique biogeography and sampling regime (Text S2) of each likely interact, making it difficult to disentangle these effects. Unsupervised learning further confirmed that each dataset exhibited unique properties (Fig. 2a), highlighting the need for caution when attempting to infer ecological variables using models or findings from another dataset (43).

### 3.2. Performance and comparison of machine learning approaches

When selecting an unsupervised or supervised approach, we find that the former has several advantages over supervised learning. Firstly, the natural overlap that can occur between soundscapes across time is more gracefully reported by unsupervised learning, whereas this often leads to a classification error during periods of temporal convergence if using supervised learning. Supervised learning is also limited by the sensitivity of classification accuracy to the training and evaluation split of the data (Fig. 3), requiring careful curation of this alongside extensive repeats to ensure confidence in performance, which can be difficult with the low spatial replication typical of marine PAM studies. Additionally, further measures can be extracted from unsupervised outputs, such as cluster diversity across recording samples from any given site, percentage overlap between cluster allocations of recordings from different sites, centroid distances in the embedding space, and more.

With regard to selecting the optimum feature extraction approach for ML, a clear difference in performance between the methods was revealed by the unsupervised learning tasks. Here, the T-CNNs produced clusters with a notably greater fidelity to true classes for every task compared to the compound index and in most tasks when compared to the P-CNN (Fig. 2C, Fig. S1), which in turn outperformed the compound index. These results reveal that feature embeddings obtained from deep learning yield improved outputs compared to handcrafted compound index embeddings.

Notably, this represents the first demonstration that transfer learning using VGGish, a network pretrained on data from an entirely different audio domain (5.2m hrs of YouTube audio), can be successfully applied to coral reef soundscapes.

For supervised classification tasks, our findings show that the difference in performance between the three embedding extraction methods on reef soundscape data was minimal. Multiple algorithms and their respective hyperparameters are available. However, it is well documented that as dataset size grows the choice of algorithm matters less (44). Our findings support this in the context of soundscapes given the broadly similar performance exhibited at the scale of the data used here.

Furthermore, datasets of this scale are easily achieved using raw soundscape data, one of the key advantages of whole soundscape analysis over bioacoustic signal analysis which requires extensive human annotation to construct labelled datasets (25). Further accuracy improvements to supervised classifiers are more likely obtainable by considering recording periods longer than the one-minute samples used here. Although short samples (e.g., one-minute) are common practice in the literature (16,24,45), a preliminary comparison using the most common classification pooled over 24 hrs in our data highlights the potential of considering longer periods (Fig. S4).

Moving forward, for guaranteed optimal performance, we recommend that users develop T-CNNs specific to their dataset for use in unsupervised approaches. However, the T-CNN approach requires significantly more expertise and computational overhead (Text S4). These overheads are greatly reduced for the P-CNN at the price of only a small performance dip in the tasks presented here. The P-CNN also represents a more standardised tool than a compound index which requires curating from an extensive set of indices and parameters for each index that currently vary from study to study (29,37). The standardised features of a P-CNN also enable comparisons of new datasets more readily, whereas a T-CNN would require retraining. We therefore recommend a P-CNN for most use cases.

### 3.3. Future directions

We present only a limited set of problem tasks, further experimentation with ML will help disentangle the relationship between soundscapes and the attributes of reefs which drive these. These experiments could be performed across ecological gradients, used to track temporal cycles, reveal patterns in functions reliant on soundscapes (46–48), or explore other novel applications. To train models that can identify relationships with raw biodiversity measures, ground-truth ecological survey data from a representative selection of sites will be essential. The impacts of different treatments (e.g., restoration, degradation events) across sites can often be compared using soundscape data alone (21,45). Prospective users should take care to follow best practices when assembling datasets by controlling for confounding variables that can impact machine learning algorithms, including instrument bias, temporal autocorrelation, careful curation of training and test sets, alongside other considerations (49–51). Future work could also aim to reap the benefits of a P-CNN while bridging the performance gap to the T-CNN by developing a generalisable reef specific P-CNN (52). Furthermore, the approaches we present here remain a ‘black-box’, meaning that the raw attributes of recordings which drive the groupings observed are unknown. Future work could improve on this by identifying the attributes (e.g., fish choruses) contributing to groupings, which could in turn be related back to the ecology.

The availability of low-cost recording technology (53,54) and rapid insightful analysis now possible using ML has the potential to greatly increase the scale of ecological assessment on coral reefs. As long as careful sampling design and evaluation protocols are in place, artificial intelligence represents a powerful tool to process this data and unlock new insights. The potential demonstrated here also has implications for other marine and terrestrial habitats where these techniques could be applied. While there remains much to discover, soundscape ecology has a lot to offer in terms of understanding, protecting and restoring coral reefs and other habitats around the world.

### 4.0 Methods

### 4.1. Datasets

The datasets used in this study were collected from three different coral reef biogeographic realms: (1) South Sulawesi, Indonesia, in the Tropical Indo-Pacific; (2) Lizard Island, Australia, in the Coral Sea; and (3) French Polynesia, in the Mid-South Tropical Pacific (Fig. 1B)(31). Each dataset was collected using SoundTrap hydrophone recorders (SoundTrap 300ST, Ocean Instruments, Auckland, NZ) which were calibrated by the manufacturer with a flat frequency response. These were suspended 0.5 m above the seabed using a sub-surface buoy. To mitigate against instrument bias, the individual recorders were frequently rotated between sites for the Indonesian and Australian datasets, whereas logistical constraints of deploying at mesophotic depths prevented this for French Polynesia (Text S2). To minimise the introduction of geophonic noise, all recordings were taken during seastates between 0-2 on the Beaufort scale and never during periods forecasted to rain, where conditions deviated from this, recordings were not taken. All recordings were taken in remote areas away from frequently trafficked boating channels and mooring points to minimize the presence of anthropogenic noise (Text S3). Further details including, maps, recording schedules and instrument rotations are available in Text S2.

### 4.2. Extracting feature embeddings

Guidance from previous literature was used to assemble a compound index. The aim was to include a broad selection of features, whilst preventing this from including unnecessary noise which can reduce the performance of machine learning algorithms. This begun with the identification of suitable acoustic indices using the following criteria: (i) previous use in the literature on marine soundscape recordings, (ii) do not require recorder calibration (precluding the use of more widely accessible recorders (53); (iii) could be computed using existing toolkits in one programming language (e.g., Python). Of the identified indices, the acoustic diversity index (ADI) and acoustic evenness (AEI) have previously been reported to strongly covary (21). Therefore, to reduce the introduction of noise only AEI was used (55) (Table S3). Seven of the eight indices were then calculated across three different frequency bands: a low-frequency (0.05–2 kHz) where fish sounds dominate (5), a medium-frequency band (2–8 kHz) where snapping shrimp sound dominates (21) and a full band (spanning 0.05–8 kHz). We excluded frequencies below 0.05 kHz from the low- and broad-frequency band recordings to remove geophonic noise and self-noise from the recording system (56). The exception was the normalised difference soundscape index (NDSI), which requires the input of two bands, for which 0.05–1 kHz and 2–5 kHz bands were used as in Williams et al., (2022). Recordings were split into 0.96 second segments (to match CNNs), totalling 62 per minute, and indices were calculated for each segment. The mean and standard deviation of these were then taken for each minute (16), providing a 44 dimension embedding which was used as the compound index to represent each one-minute recording. All processing was performed in Python (v3.7) using the scikit-maad package (v1.3)(57).

For the pre-trained convolutional neural network (P-CNN), we selected VGGish due to its successful application in previous work on terrestrial soundscape data (24,58). VGGish was trained using the YouTube-100M dataset, a diverse set of YouTube audio clips totalling 5.2m hrs in length, producing a highly generalisable audio and embedding extractor (28). It uses a version of the Visual Geometry Group object recognition architecture that was adapted for audio input (59). The P-CNN pre-processing first down-samples recordings to 16 kHz, and divides them into non-overlapping 960 ms audio frames, processed through a Short-time Fourier Transform with 25 ms windows every 10 ms. These are integrated into 64 mel-spaced frequency bins which are log-transformed to produce a 96×64 bin log-mel spectrogram which can be passed to the network. Further details are available in Hershey et al., (2017). This P-CNN can be set to output the 128-dimension feature set from the penultimate layer in place of the classification head. We averaged these feature values from each one-minute recording to produce a single embedding processing was performed in Python (v3.7).

To ensure comparability with the P-CNN, we trained VGGish on each respective reef soundscape dataset and task to produce the T-CNNs. The pre-processing protocol used for the P-CNN was replicated to produce 960 ms log-mel spectrogram samples. The order of these samples was shuffled and minibatches of 32 samples used for network for training. Processing was performed using Tensorflow (v1.15) and scikit-learn (v0.22) in Python (v3.7).

### 4.3. Unsupervised clustering

Unsupervised clustering was used to reveal inherent structures and patterns in the data without relying on pre-labelled classes. Embeddings from the compound index and P-CNN were extracted from recordings. To generate embeddings using the T-CNNs, the VGGish CNN was trained for 50 epochs on each dataset to predict which site recordings originated from within the respective dataset (see section 4.4 for more detail). The final layer of the trained models were then removed to produce three networks comparable to the P-CNN, except these were now trained on reef soundscape recordings from their respective datasets. These T-CNNs were then used as embedding extractors on all recordings from their respective datasets in the same manner as the P-CNN. T-CNNs trained to classify from which site recordings originated were used for both the site and habitat classification tasks given that the site of origin, but not habitat type, should be known for any dataset.

To improve clustering, dimensionality reduction was performed on each embedding using uniform manifold approximation (UMAP) via its associated Python package (v0.5.3)(32). To determine which of the three embedding extraction methods was most proficient at producing UMAP embeddings that represented known properties of the data, a qualitative assessment was first performed using UMAP visualisation in two dimensions. For all six tasks performed by each method, a plot was produced where points were labelled with their known class and the fidelity of groupings to their true class compared between each method.

A quantitative assessment was also undertaken using affinity propagation clustering (60) on each UMAP embedding. This algorithm was selected as there is no requirement to predetermine the number of classes, instead the affinity propagation algorithm identifies this, simulating analysis with an entirely unlabelled dataset where number of classes is unknown. For a given method and task, the cluster to which each recording was assigned was entered into a contingency table against its known class. Models most proficient at clustering recordings from the same class, while excluding recordings from other classes, generated more discrete clusters. Those less proficient assigned recordings more randomly across clusters. A chi-squared test was performed on contingency tables to assess this, with a higher score indicating models with a higher fidelity to true classes. Processing was performed using scikit-learn (v0.22) in Python (v3.7).

### 4.4. Supervised classifiers

All three ML methods were set two classification tasks (habitat and individual site identity) from each of the three soundscape datasets. The habitat classification tasks were as follows: high or low coral cover sites for the Indonesian dataset, high or low fish diversity sites for the Australian dataset, and shallow or mesophotic sites for the French Polynesian dataset. For each task, recordings were split into training, validation and test sets, where 66-75% of the data was used for training and the remaining data was evenly split for validation and testing of each task (see Text S2 for specific details for each dataset and task). Validation and test accuracies were carefully monitored to ensure clear over and underfitting did not occur.

For the compound index and P-CNN embeddings, random forest ML classifiers were trained for each classification task. Random forests were selected due to their previous successful implementation in the soundscape literature (16,24), robustness to overfitting, and, low computational costs which enable calculation on personal computing devices; a likely consideration for future users (Text S4). For each task, fifty random forest classifiers were trained and then used for inference on the validation data. The instance which reported the highest validation accuracy was used for inference on the test data, and the accuracy of this reported. As classes were well balanced within each task, raw accuracy (the proportion of one-minute recordings correctly classified) was used as the performance metric. For each task, this process was repeated multiple times using different splits of the data into training, validation and test sets as initial trials showed the split impacted performance of the three embedding approaches differently. One-hundred of these repeats were performed for each task, except for the Australian and French Polynesian habitat classification tasks, where only 32 were used as this was the maximum number of possible combinations which enabled the exclusion of entire sites from the training data (Text S2). Analysis was performed using scikit-learn (v0.22) in Python (v3.7).

For the T-CNNs, these were trained on the reef soundscape dataset for each individual task. The number of output nodes was adjusted to match the number of target classes in each task. The T-CNN was trained for 50 epochs, with inference on the validation data every epoch. The epoch which reported the highest validation accuracy was then used to inference on the test data. The most common class prediction across all 960 ms segments in a one-minute recording was used as the class prediction for each minute in the test data. This process was repeated 100 times using the same training, validation and test combinations as the compound index and pretrained T-CNN. This workflow was performed using Tensorflow (v1.15) in Python (v3.7).

Analysis of variance (ANOVA) tests were used to determine whether any significant difference between the accuracy of each embedding approach was present across all repeats performed for each task. Where significant differences were detected (p<0.05), post hoc Tukey tests were used to determine which methods differed; performed using the scipy library (v1.73) in Python (v3.7).

## Acknowledgements

For providing the Indonesian dataset the authors would like to thank Mars Global; Hasanuddin University; the Department of Marine Affairs and Fisheries of the Province of South Sulawesi; the Government Offices of the Kabupaten of Pangkep, Pulau Bontosua and Pulau Badi; and the communities of Pulau Bontosua and Pulau Badi for their support. For the Australian dataset the authors would like to acknowledge the Dingaal Aboriginal people (original land owners of Lizard Island) and thank staff from Australian Museum Lizard Island Research Station. For supporting the collection of the French Polynesian dataset the authors would like to thank staff from the Centre de Recherches Insulaires et Observatoire de l’Environnement (CRIOBE). This work was funded by the following grants: Fisheries Society of the British Isles PhD studentship award to B.Ws; Natural Environment Research Council Research Grant to S.D.S (NE/ P001572/1); Natural Environment Research Council–Australian Institute of Marine Science CASE GW4+ Studentship (NE/L002434/1) and a Royal Commission for the Exhibition of 1851 Research Fellowship to T.A.C.L; Swiss National Science Foundation Early Postdoc Mobility fellowship to L.C (P2SKP3-181384); Agence National de la Recherche to Glenn Almany, S.C.M and R.B (ANR-14-CE02-0005-01/Stay or Go); Haut-Commissariat de la République en Polynésie Française to S.C.M (HC/3041/DIE/BPT/); Pacific Funds (BLEACH & ALAN) to S.C.M; Millenium Nucleus for the Ecology and Conservation of Temperate Mesophotic Reef Ecosystem (NUTME), Chile to R.B.

## Data availability statement

We share the full datasets of raw recordings used in this study, totalling >340 hrs. Due to their size, the data is split across three Zenodo repositories. For specific details beyond those shared in this manuscript, please contact the corresponding author (BW). The raw Indonesian audio alongside feature embeddings extracted from the full datasets, T-CNN predictions and a link to sample audio are available from: https://doi.org/10.5281/zenodo.11106482.

The remaining raw audio data can be found in these repositories:

Australia: https://doi.org/10.5281/zenodo.10533066

French Polynesia: https://doi.org/10.5281/zenodo.10539938

## Code reporting

A full tutorial on running our recommended method, a pretrained network with unsupervised learning, is available alongside access to sample data. This can be run entirely from a web browser using Google Colab and the sample data stored on Google Drive: https://github.com/BenUCL/Reef-acoustics-and-AI/tree/main.

The remaining code used to complete the study is available from the same GitHub repository and can also be accessed from: https://doi.org/10.5281/zenodo.11106482.

## Notes

https://doi.org/10.5281/zenodo.11106482

https://doi.org/10.5281/zenodo.10533066

https://doi.org/10.5281/zenodo.10539938

https://github.com/BenUCL/Reef-acoustics-and-AI

### Competing Interest Statement

The authors have declared no competing interest.

### Summary of Updates

Fixed a spelling error in author name

