## Supplementary 1 for "Unlocking the soundscape of coral reefs with artificial intelligence: pretrained networks and unsupervised learning win out"

### Supplementary Texts

#### Text S1: Interactive UMAP plots

Interactive UMAP plots can be accessed via Supp. 2: https://doi.org/10.5281/zenodo.11106482.

Here a zip file can be downloaded which contains HTML files for an interactive UMAP plot from each dataset. Once opened, users can hover their cursor over the individual points to reveal temporal metadata about each point and the site of origin. Points are coloured by site. Recordings from similar times periods show a clear pattern of grouping together. Further exploration also reveals other temporal insights. For example, on the French Polynesian plot, the 'bridges' between primary clusters for each site are dominated by recordings taken during crepuscular periods where 'hr of day' is 5-7am or 17-19pm. In the Indonesian plot, an area of overlap between site A and C is visible during the new moon period at 5pm and 5am respectively, showing the soundscapes converged during this period. Other patterns can likely be revealed using this tool.

#### Text S2: Recording schedules and train, validation, test divisions

Recordings were taken in multiple discrete blocks. Across each dataset, hydrophone recorders were frequently rotated between sites for a new recording block to mitigate against the introduction of instrument bias. For habitat classification tasks, entire sites were excluded from the training data for the Australian and French Polynesian dataset so that unseen sites were evaluated upon. The Indonesian dataset only had two sites in each category, so entire recording blocks were excluded for this. For site classification tasks, entire recording blocks were excluded from the training data and used to evaluate upon. This was to mitigate against temporal autocorrelation, meaning a classifier would not be able to train upon recording periods immediately adjacent to those in the evaluation set. Further details on how this was implemented for each dataset are below.

**Indonesia**

The Indonesian dataset was collected as part of the monitoring programme of the Mars Coral Reef Restoration Project (www.buildingcoral.com). This monitoring programme included recordings collected from four study sites across two reefs (Fig. S5b); two high coral cover reefs (Fig. S5c-d) and two low coral cover reefs (Fig. S5c). All sites were at a depth between 2–3.3m at low tide. One-hour blocks of soundscape recordings were taken at each site for five days either side of the full moon (26 August 2018) and three days either side of the following new moon (10 September 2018) during daylight (09:00–15:00), twilight (05:30–06:30 and 17:30–18:30) and night-time (23:30–00:30) periods. These were made in a counterbalanced blocking design, such that there was a similar number of recordings taken from each site, comprising an approximately even spread of time points and lunar phases (see Lamont et al., (2021) for more detail). Recordings were split into one-minute segments, providing a total of 3,335 one-minute recordings.

For the Indonesian habitat and site level classification training and test divisions, 57 one-hour recording blocks were available, split approximately evenly across the four sites (Fig. A)(see Lamont et al., 2021 for further details). For both tasks, eight blocks were selected for the test set and eight for the validation set (Fig. A). The remaining 41 blocks were used for the training set.

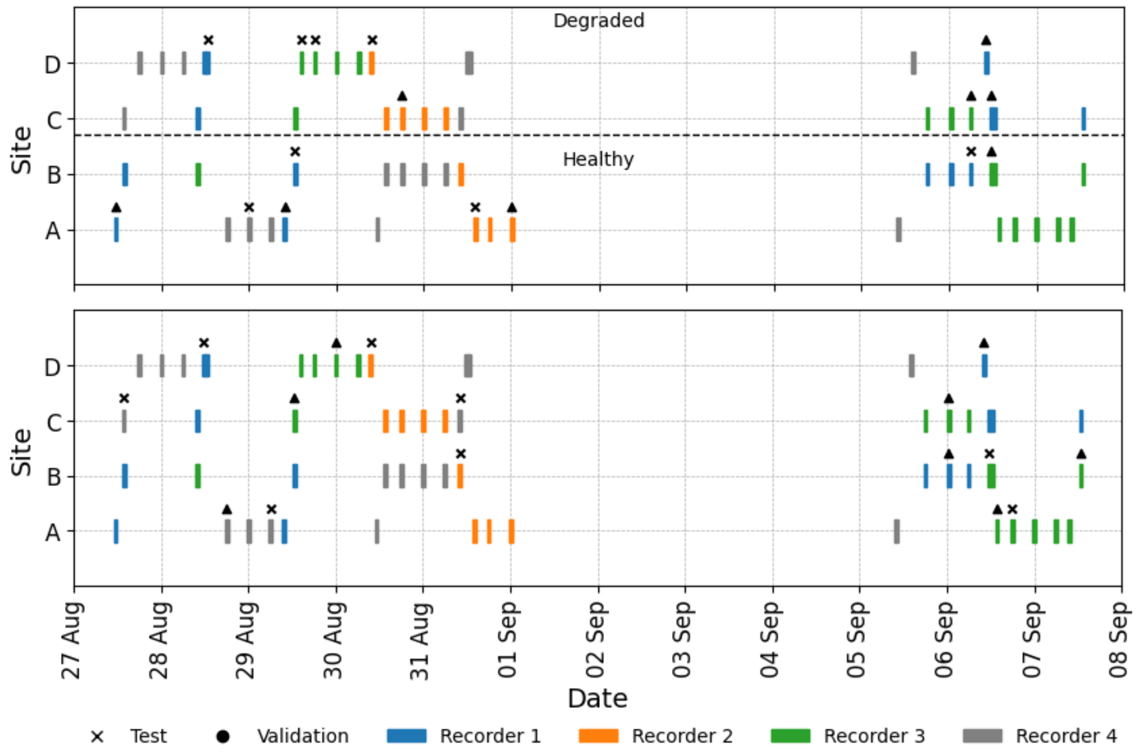


**Fig. A.** Plots depicting the recording schedule and instrument rotation followed for the Indonesian dataset. Gridlines represent midnight. Recordings were taken after the full moon (Aug 26$^{th}$) and before the new moon (Sep 10$^{th}$). (Top) For the habitat classification task, four blocks were randomly selected from each of the two classes as validation data, and another four as test data, remaining blocks were used as training data. A new combination was selected for each of the 100 repeats performed. The split for one example repeat is depicted, where symbols above blocks indicate those which were used for the test or validation in this repeat. The same combinations were conserved across compound index, P-CNN and T-CNN training. (Bottom) Site identification task. Identical to the habitat task except two blocks were randomly selected from each site of the four sites as validation data, and another four as test data.

### Australia

The Australian dataset consisted of recordings from 12 randomly selected sites (A–L) located around Lizard Island on the Great Barrier Reef; sites were separated by at least 500 m and constituted >250 x 5 m of contiguous reef (Fig. S5e). All sites were at a depth between 2–3m at low tide. Six recording blocks of 19–24 hr duration were taken at each site between 23 October to 14 December 2018; blocks from each site were spaced 9 days (SE: ±3.49) apart on average (Text S2). Recordings were not made during the Nov 11-20$^{th}$ period due to elevated sea states which could introduce geophonic noise. Recordings were subsampled to take one-minute clips separated by nine minutes of unused recording, providing 8,127 one-minute recordings in total. Fish community biomass and species richness data were collected along three transects at each site during this period following methods from Richardson et al., (2018). Of the 12 sites, the four with the highest and four with the lowest species richness values were identified and split into two classes, totalling 6289 one-minute recordings after excluding the other four sites (Fig. S6). The sites from these two classes were also found to have non-overlapping fish biomass values and were therefore split into high and low fish diversity classes.

For the habitat level classification training and test divisions, 48 19-24 hr recordings blocks were available, split evenly across the four high and four low fish diversity sites (Fig. B). For the site level classification training and test divisions all 12 sites were included, providing 72 one-day recordings blocks split evenly across these sites (Fig. B).

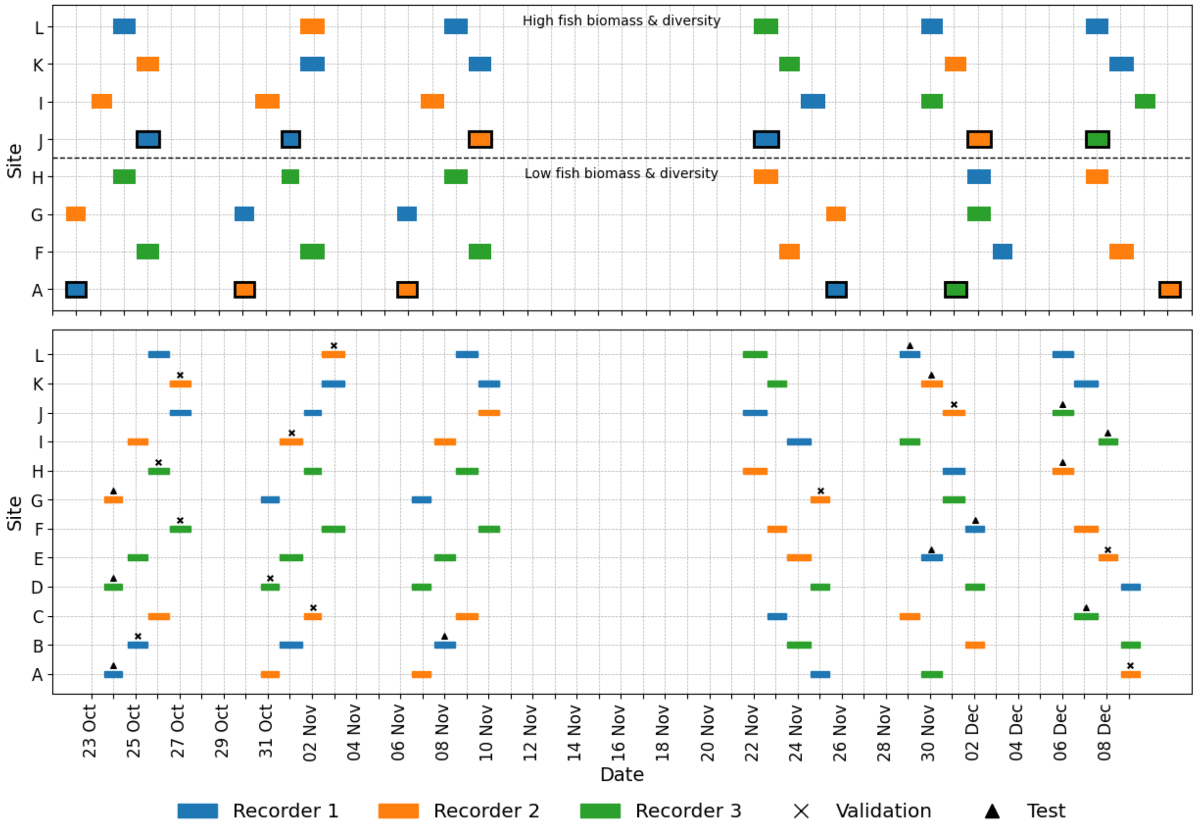


**Fig. B.** Plots depicting the recording schedule and instrument rotation followed for the Australian dataset. Gridlines represent midnight. (Top) For the habitat classification task, one entire site was taken from both classes and all recording blocks from these two sites aggregated and randomly split in two whilst maintaining a balanced class ratio to provide validation and testing data. This was then repeated, with the validation and test set swapped to provide another repeat. This was repeated for every combination of the eight sites, maintaining one high and one low fish diversity site in the validation and training set each time, producing 32 replicates in total. Here, one example repeat with its validation and test set is depicted by the blocks with a black outline. (Bottom) For the site classification task, one recording block was randomly selected from each site as the validation data, and another block for the test data. Remaining blocks were used as training data. A new combination was selected for each of the 100 repeats performed for this task. One example repeat is depicted, where shapes above blocks indicate those which were used for the test or validation in this repeat. The same combinations were conserved across compound index, P-CNN and T-CNN training.

**French Polynesia**

The French Polynesian dataset was collected in February and March 2021 at four sites, consisting of reefs around the islands of Mo'orea, Tikehau and Tahiti (Fig. S5f-h). At each site, soundscape recordings were taken at a shallow (10–15 m depth) and mesophotic (55–65 m depth) site, totalling four shallow sites (A–D) and four mesophotic sites (W–Z). Recordings were taken simultaneously for each shallow–mesophotic pair for a continuous period of 89–97 hrs (Text S2). Recordings were subsampled to take one-minute clips separated by four minutes of unused recording, providing 8,975 one-minute recordings in total.

For training and test division, 89-97 hr blocks recording blocks were taken at each of the eight sites (Fig. C). For the habitat classification task, entire sites were set aside as validation and testing data. For the site classification task, contiguous 24 hr recording blocks were excluded and set aside as validation and testing data. Unlike the other datasets, the recorders could not be frequently rotated in the French Polynesian dataset due to the logistical constraints of using technical divers to deploy the hydrophones at mesophotic depths.

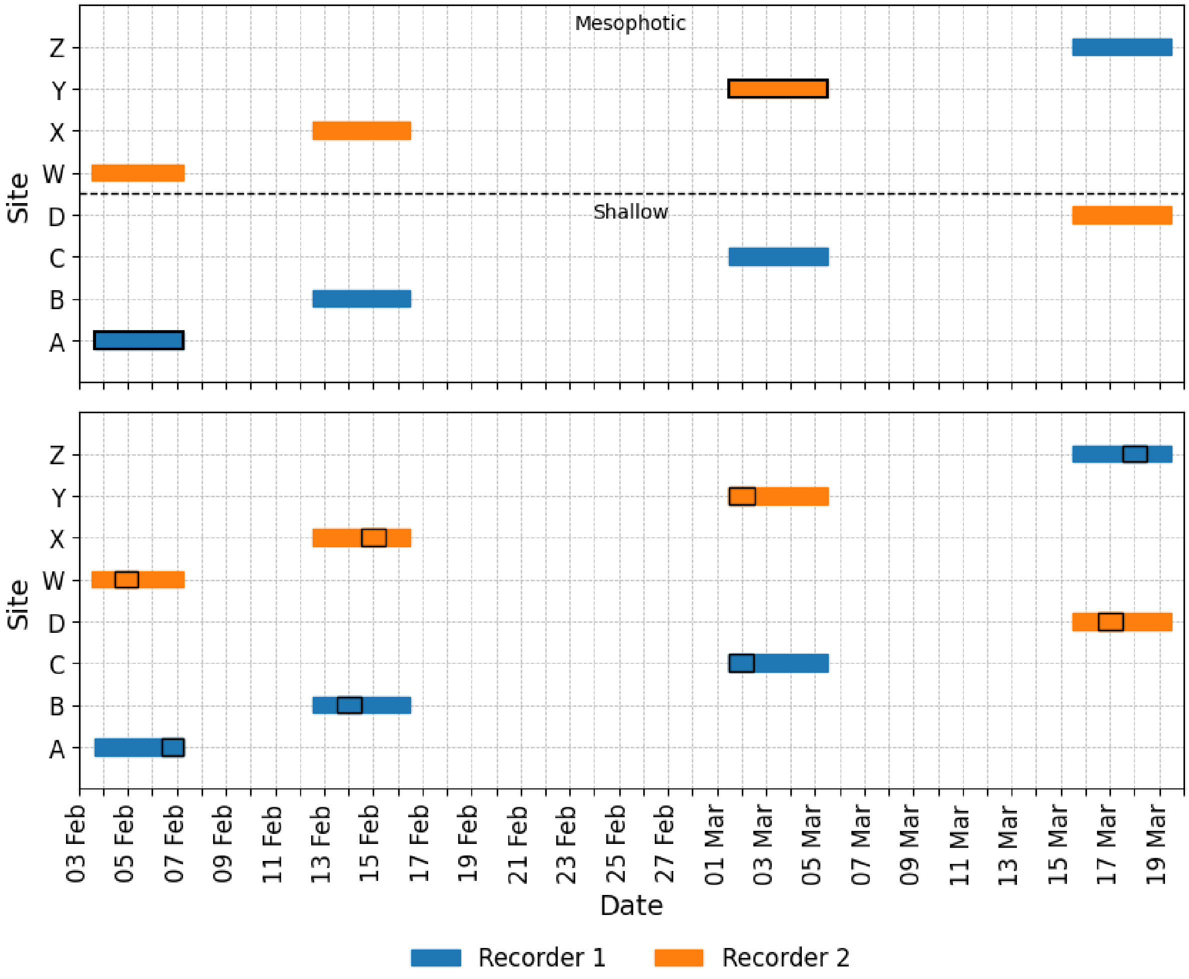


**Fig. C.** Plots depicting the recording schedule and instrument rotation followed for the French Polynesian dataset. Gridlines represent midnight (Top) For the habitat classification task, one site was taken from both classes. All one-minute recordings were randomly split in two whilst maintaining a balanced class ratio to provide validation and testing data. This was then repeated, with the validation and test set swapped. This was then repeated for every combination of the eight sites, maintaining one shallow and one mesophotic site in the validation and training set each time, producing 32 replicates in total. Here, the validation and test set for one example repeat is indicated by the blocks with a black outline. (Bottom) For the site classification task, recording blocks were split into 24 hr periods and one period was randomly selected. This selected period was randomly split in two to provide validation and test sets. Remaining periods were used as training data. A new combination was selected for each of the 100 repeats performed for this task, one example combination is depicted by periods with black outlines representing the data divided into validation and testing sets. The same combinations were conserved across compound index, P-CNN and T-CNN training.

### Text S3: Motorboat noise checks

If extensive boat noise were present in some sites or habitat types but not others, this could enable models to learn features relevant to boat noise and not the biophony. Boat noise is documented to interact with elements of the ecology on reefs[1–3] which may make its presence in the soundscape an important indicator. Future studies should consider the removal or inclusion of boat noise carefully.

To check for excessive boat noise, two-hundred one-minute recordings were sampled from each dataset, evenly distributed across sites but otherwise at random. The first 30-seconds of each recording were checked for boat noise by BW, totalling 5hrs of data. None of the samples from the Indonesian dataset contained boat noise. Two samples from both the Australian and French Polynesian datasets contained boat noise, from site D and G, and, site W and A respectively. This indicates approximately 1% of the data contained boat noise. If boat noise is not evenly distributed across sites the models would still have to learn features from the biophony for 99% of samples.

### Text S4: Computational resources

The T-CNN was significantly more computationally intensive than the other two methods. The Indonesian dataset represented the smallest of the three, with 3,335 one-minute recordings. Training each repeat of the T-CNN site classifier took ~ 2hrs to extract and save log-mel spectrograms from recordings using a CPU, then ~1 hr to train on a GPU (NVIDIA A100), an often prohibitively expensive piece of equipment to purchase and host, or access via cloud computing, that also requires additional technical expertise to use. Training with a CPU available on a standard personal computing device was unable to complete even one of the 50 training epochs in a 24 hr period. Conversely, the computational resources required to run the compound index and P-CNN are much more accessible, taking 84 and 96 minutes respectively to extract embeddings from all recordings in the Indonesian dataset. Following embedding extraction, the execution time of downstream processing using random forest classifiers and unsupervised learning was negligible on a CPU.

### **Additional result figures:**

Including: Fig. S1 UMAP plots; Fig. S2 confusion matrices; Fig. S3 use of individual acoustic indices; Fig. S4 classification using 24h hr periods.

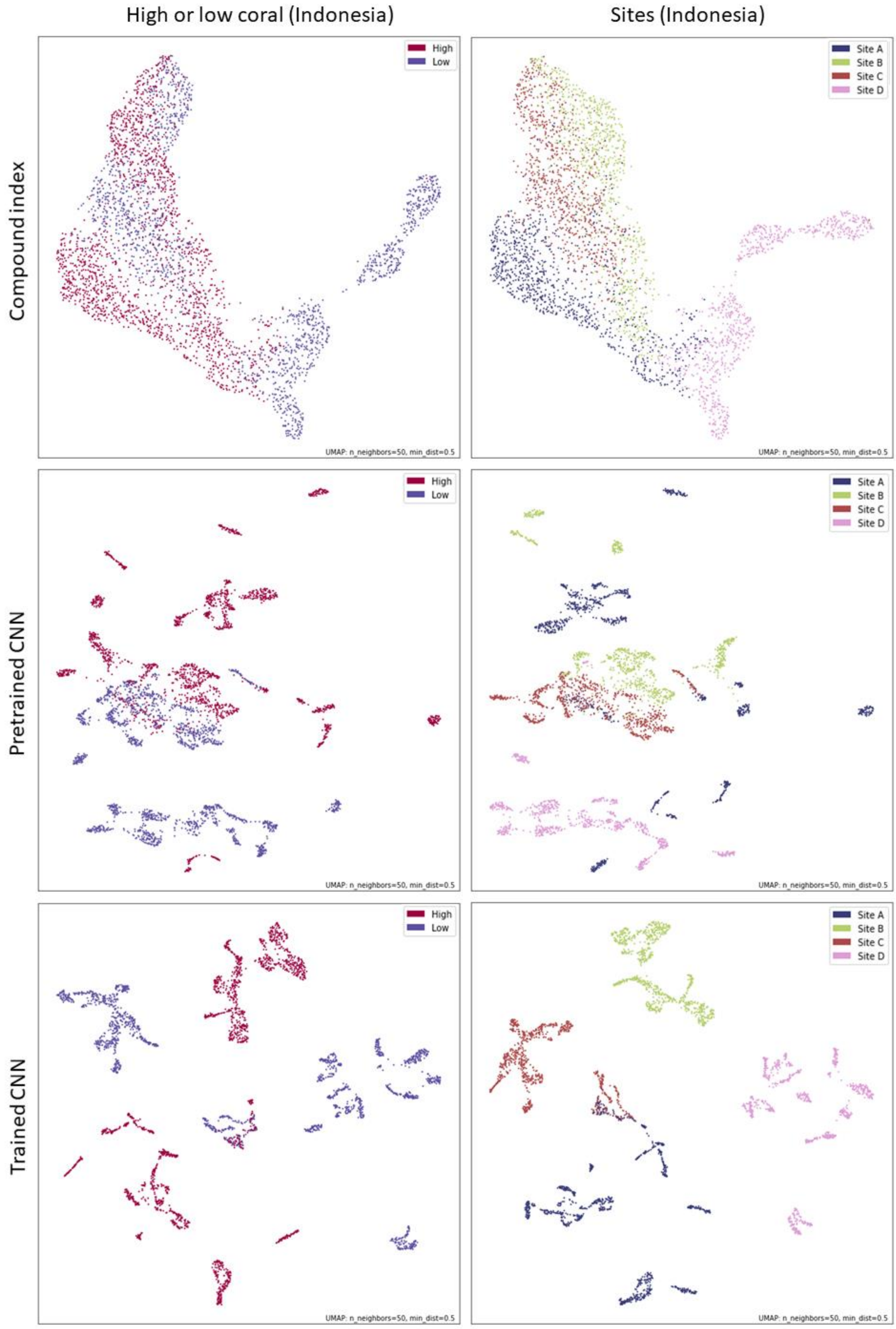

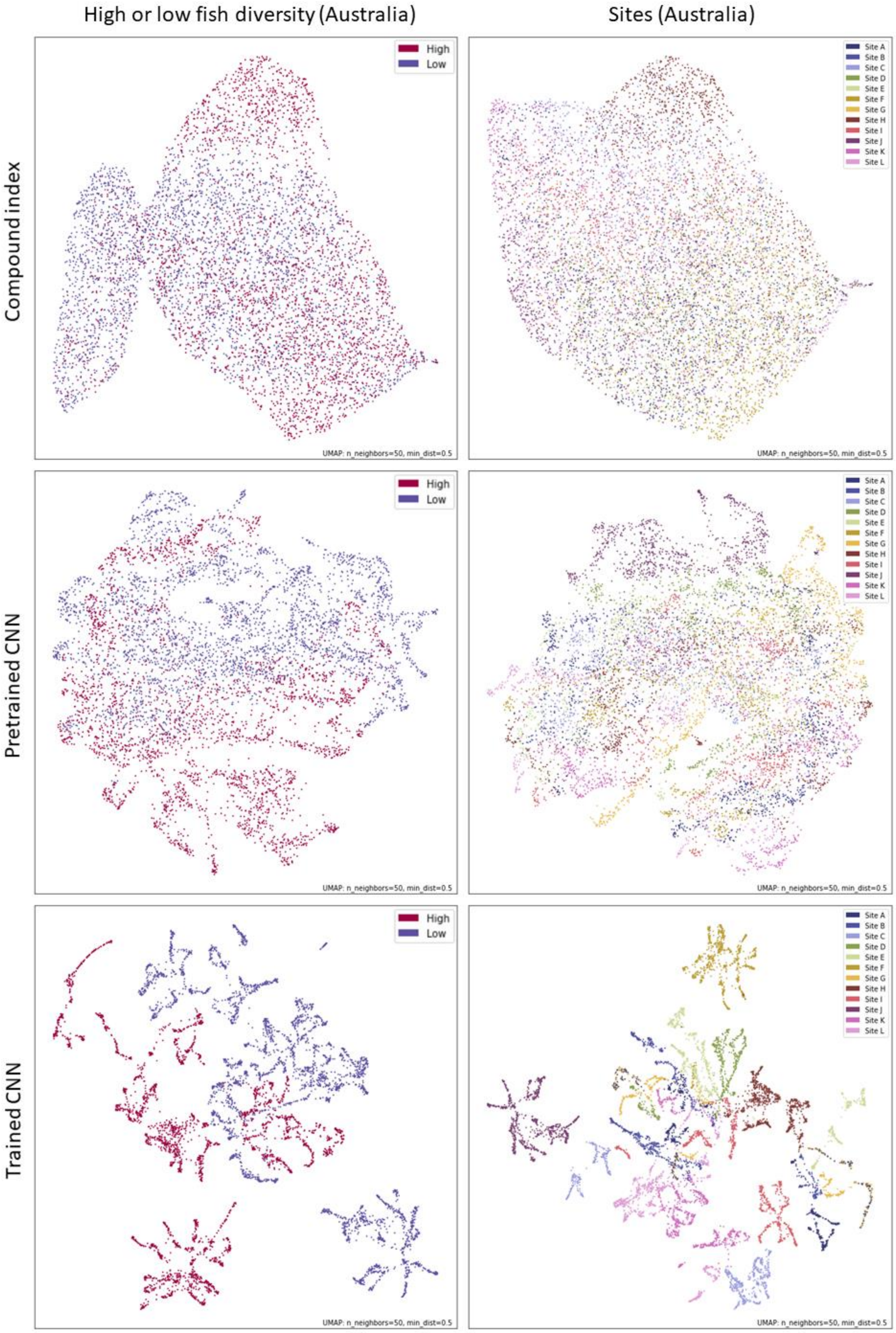
High or low fish diversity (Australia)
Sites (Australia)
Compound index
Pretrained CNN
Trained CNN
High
Low
Site A
Site B
Site C
Site D
Site E
Site F
Site G
Site H
Site I
Site J
Site K
Site L
UMAP: n_neighbors=50, min_dist=0.5

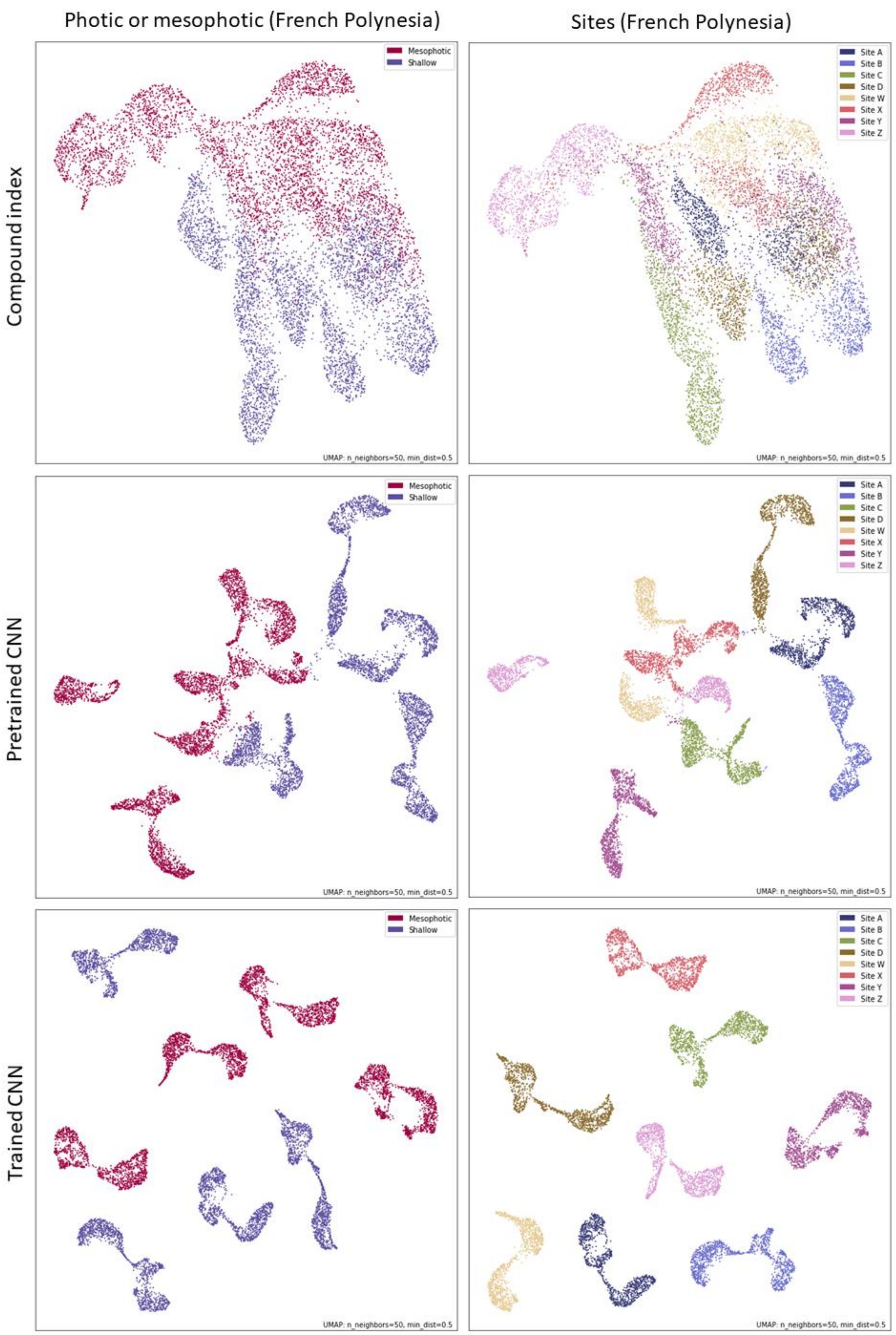


**Fig. S1.** Uniform manifold approximation (UMAP) plots used to represent the compound index, pretrained CNN and trained CNN embeddings in two dimensional space. Individual points represent a one-minute recording. Plots were produced for each of the Indonesian, Australian and French Polynesian datasets and are labelled with colours corresponding to either site or habitat class.

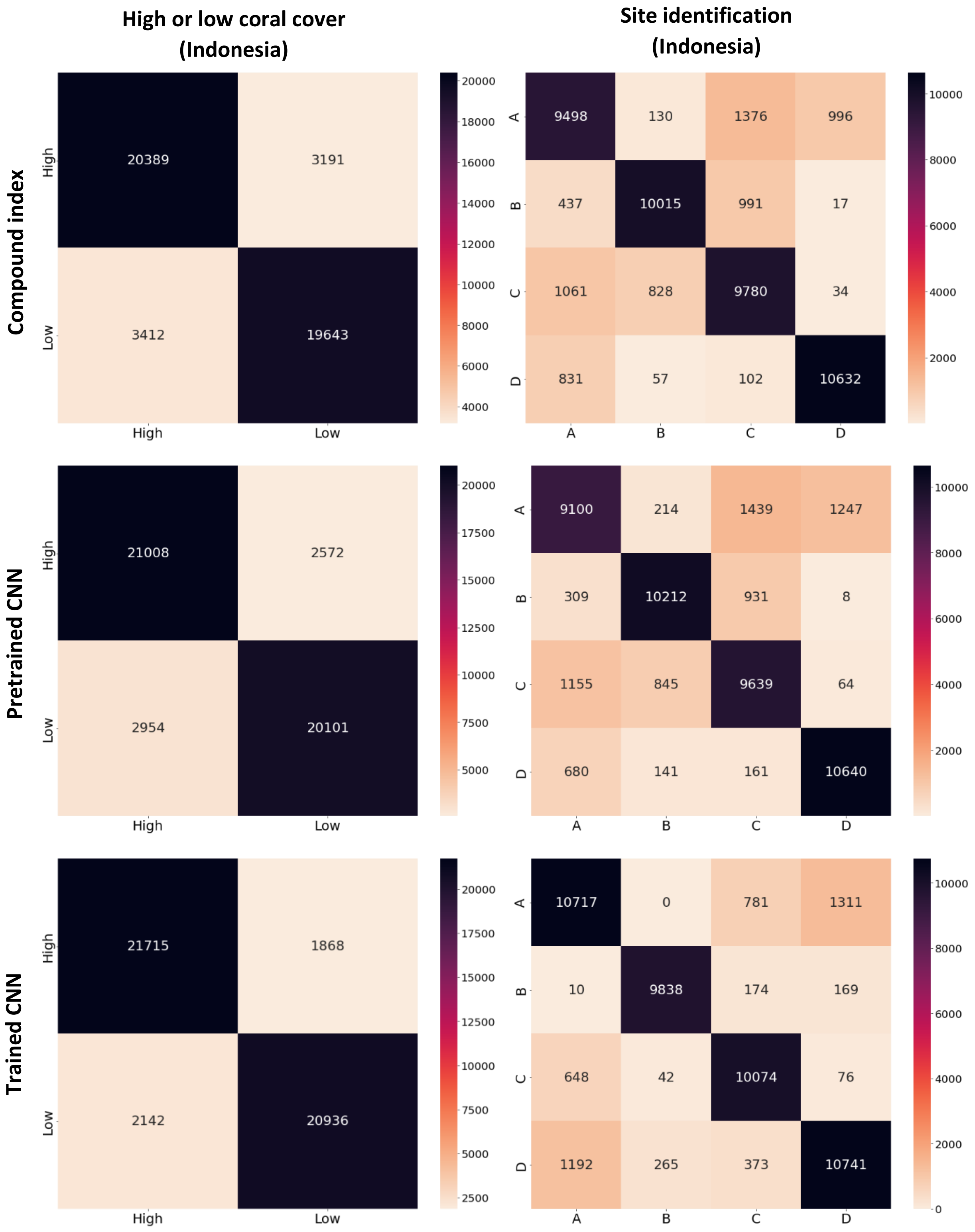
High or low coral cover (Indonesia)
Site identification (Indonesia)
Compound index
Pretrained CNN
Trained CNN
High
Low
A
B
C
D
20389
3191
3412
19643
9498
130
1376
996
437
10015
991
17
1061
828
9780
34
831
57
102
10632
21008
2572
2954
20101
9100
214
1439
1247
309
10212
931
8
1155
845
9639
64
680
141
161
10640
21715
1868
2142
20936
10717
0
781
1311
10
9838
174
169
648
42
10074
76
1192
265
373
10741

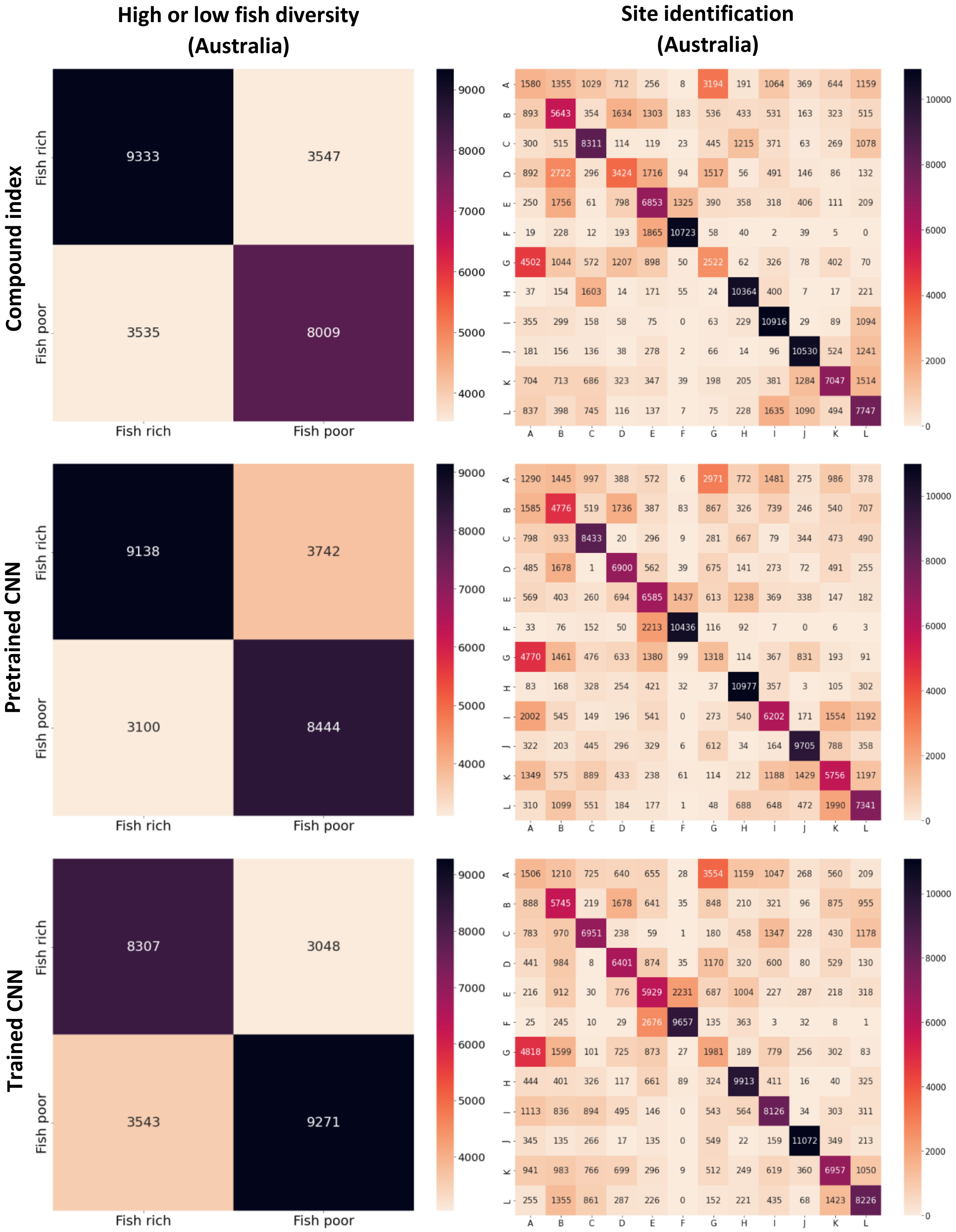
High or low fish diversity (Australia)
Site identification (Australia)
Compound index
Pretrained CNN
Trained CNN
Fish rich
Fish poor
9333 3547
3535 8009
9138 3742
3100 8444
8307 3048
3543 9271
9000 8000 7000 6000 5000 4000
10000 8000 6000 4000 2000 0
A B C D E F G H I J K L
1580 1355 1029 712 256 8 3194 191 1064 369 644 1159
893 5643 354 1634 1303 183 536 433 531 163 323 515
300 515 8311 114 119 23 445 1215 371 63 269 1078
892 2722 296 3424 1716 94 1517 56 491 146 86 132
250 1756 61 798 6853 1325 390 358 318 406 111 209
19 228 12 193 1865 10723 58 40 2 39 5 0
4502 1044 572 1207 898 50 2522 62 326 78 402 70
37 154 1603 14 171 55 24 10364 400 7 17 221
355 299 158 58 75 0 63 229 10916 29 89 1094
181 156 136 38 278 2 66 14 96 10530 524 1241
704 713 686 323 347 39 198 205 381 1284 7047 1514
837 398 745 116 137 7 75 228 1635 1090 494 7747
1290 1445 997 388 572 6 2971 772 1481 275 986 378
1585 4776 519 1736 387 83 867 326 739 246 540 707
798 933 8433 20 296 9 281 667 79 344 473 490
485 1678 1 6900 562 39 675 141 273 72 491 255
569 403 260 694 6585 1437 613 1238 369 338 147 182
33 76 152 50 2213 10436 116 92 7 0 6 3
4770 1461 476 633 1380 99 1318 114 367 831 193 91
83 168 328 254 421 32 37 10977 357 3 105 302
2002 545 149 196 541 0 273 540 6202 171 1554 1192
322 203 445 296 329 6 612 34 164 9705 788 358
1349 575 889 433 238 61 114 212 1188 1429 5756 1197
310 1099 551 184 177 1 48 688 648 472 1990 7341
1506 1210 725 640 655 28 3554 1159 1047 268 560 209
888 5745 219 1678 641 35 848 210 321 96 875 955
783 970 6951 238 59 1 180 458 1347 228 430 1178
441 984 8 6401 874 35 1170 320 600 80 529 130
216 912 30 776 5929 2231 687 1004 227 287 218 318
25 245 10 29 2676 9657 135 363 3 32 8 1
4818 1599 101 725 873 27 1981 189 779 256 302 83
444 401 326 117 661 89 324 9913 411 16 40 325
1113 836 894 495 146 0 543 564 8126 34 303 311
345 135 266 17 135 0 549 22 159 11072 349 213
941 983 766 699 296 9 512 249 619 360 6957 1050
255 1355 861 287 226 0 152 221 435 68 1423 8226

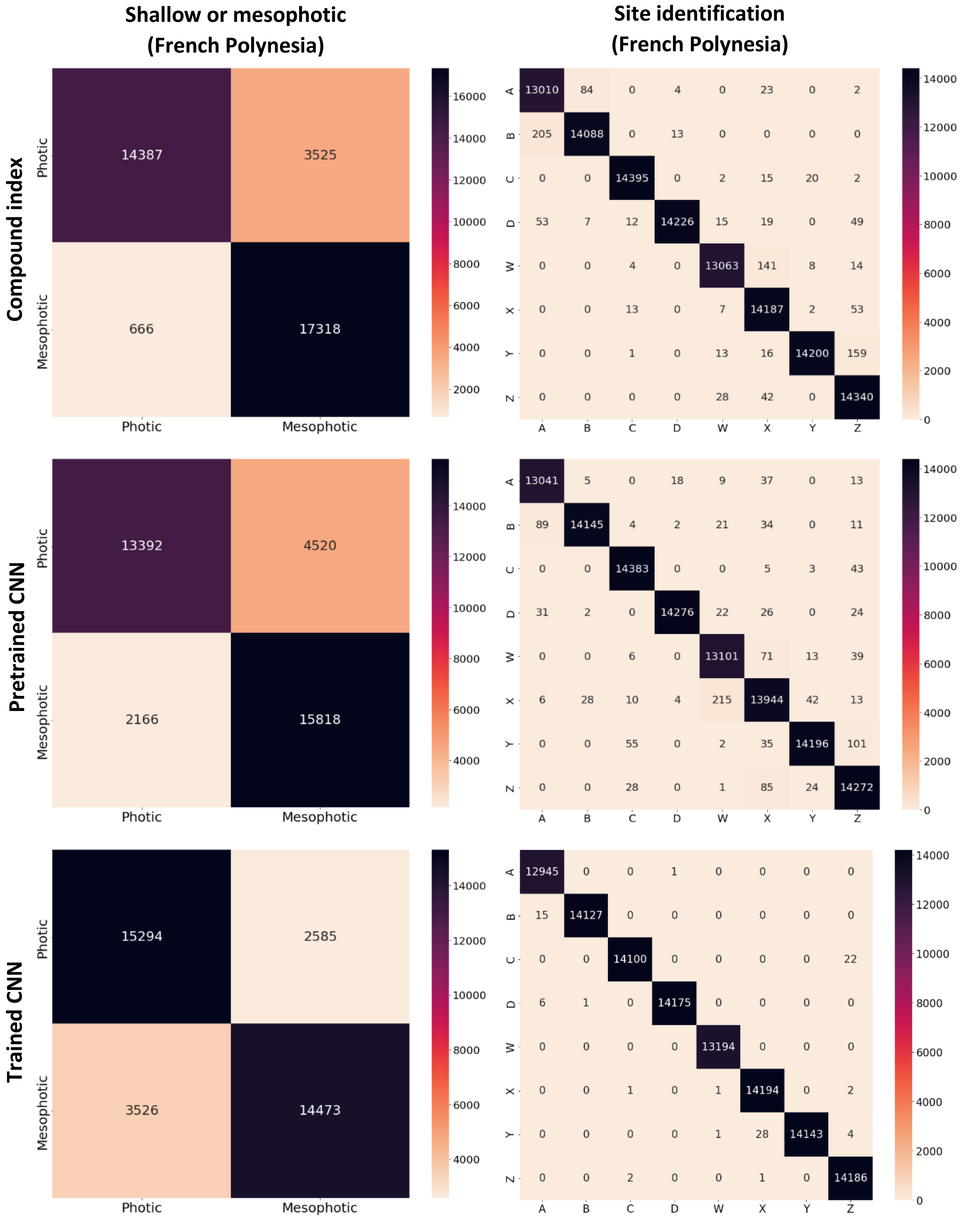


**Fig. S2.** Confusion matrices pooled across all repeats of trained classifiers for each method and task using the Compound index, Pretrained CNN and Trained CNN on the Indonesian, Australian and French Polynesian datasets. True classes are displayed along the x-axis with predicted classes across the y-axis.

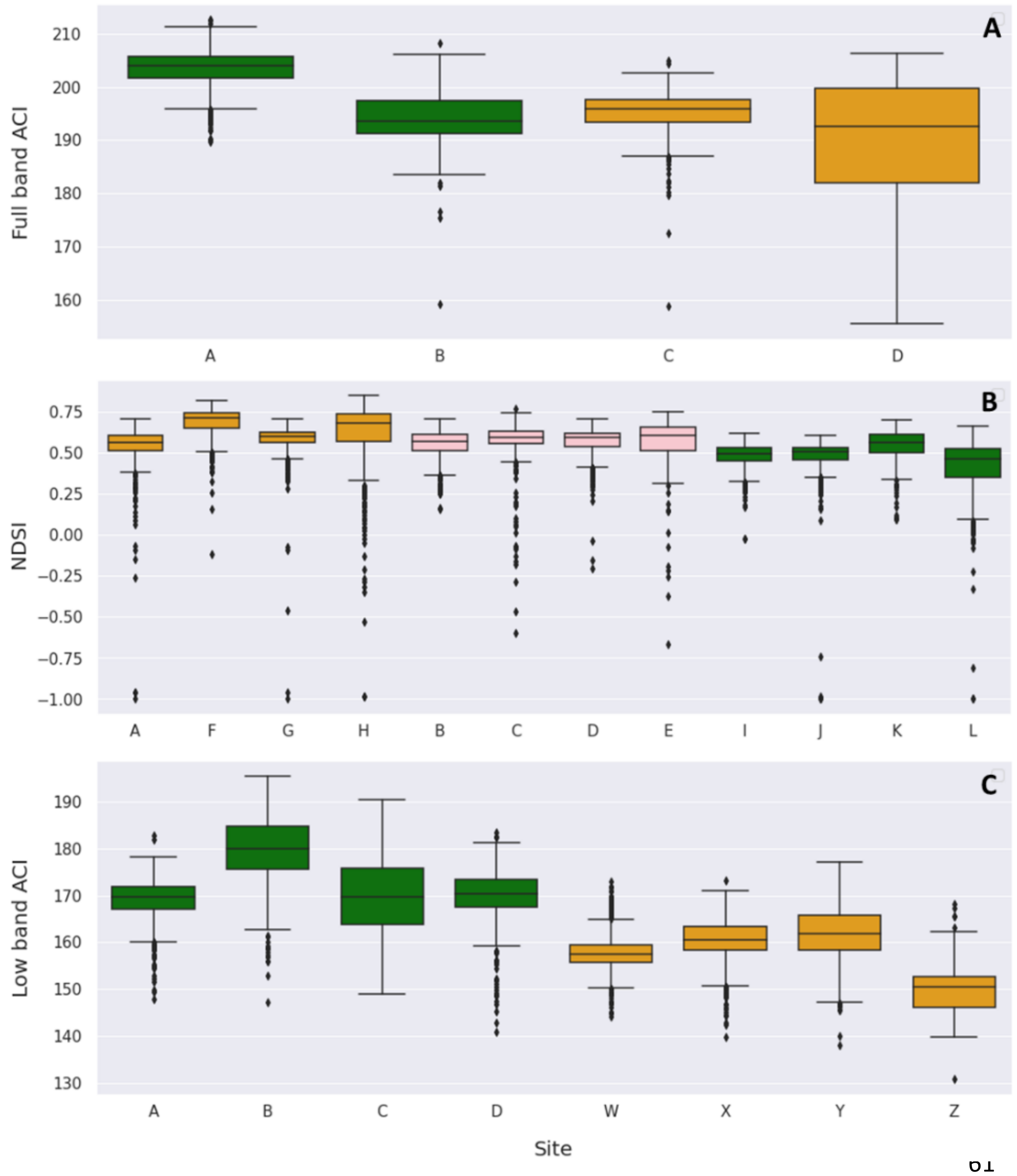


**Fig. S3.** Boxplots of individual acoustic index values for sites from the **(A)** Indonesian, **(B)** Australian and **(C)** French Polynesian datasets. Green boxes indicate high coral cover, high fish diversity and shallow reef classes for the Indonesian, Australian and French Polynesian dataset respectively, with orange indicating the opposing class, and, pink indicating the four sites excluded from habitat category task for the Australian dataset. The index with the highest significant difference between habitat classes reported for each respective dataset was selected for plotting. These were the full band acoustic complexity Index (ACI), normalised difference soundscape index (NDSI), and low band acoustic complexity index (ACI) respectively. Boxes and their bars represent the 25th, 50th and 75th quartile. The overlap of index values across sites prevents the classification of individual sites using this approach.

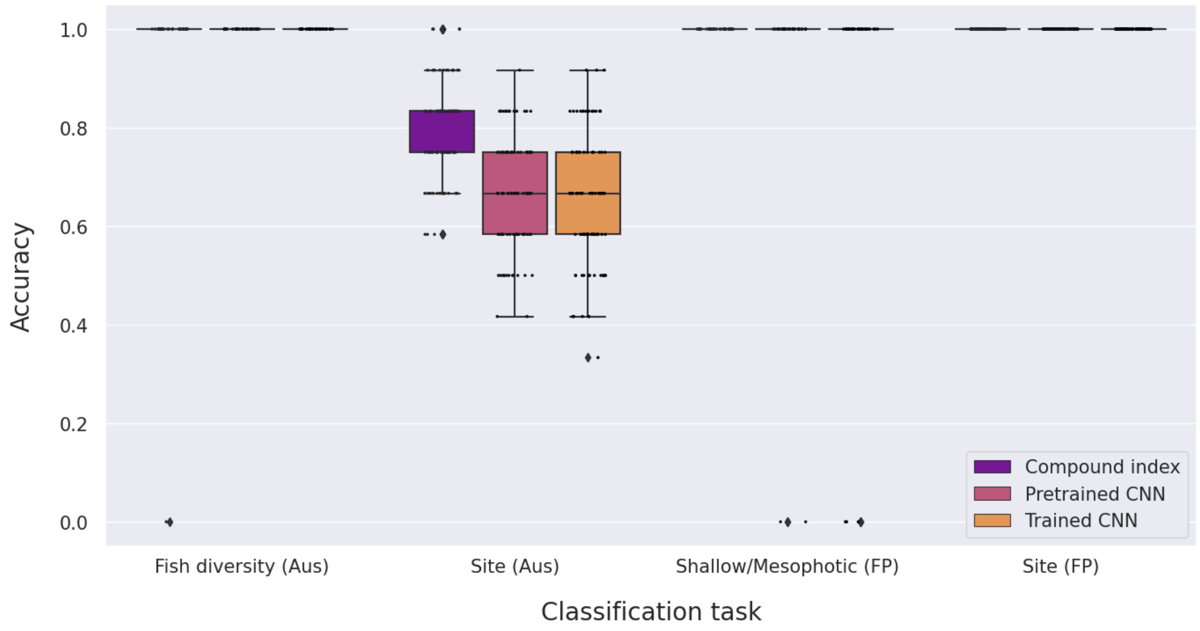


**Fig. S4.** A pilot experiment was performed which took the most common class prediction across all one-minute recordings within a 24 hr block for the Australian (Aus) and French Polynesian (FP) datasets (recordings blocks were only 1 hr for the Indonesian dataset). These results show grouping samples over longer time periods improved accuracy of all three methods. However, this reduces the sample count in the validation and test data for each repeat of the model to one, leading to accuracies of 1 or 0 for binary tasks. Future experiments may benefit from improved techniques which can consider longer periods.

### Additional methods figures

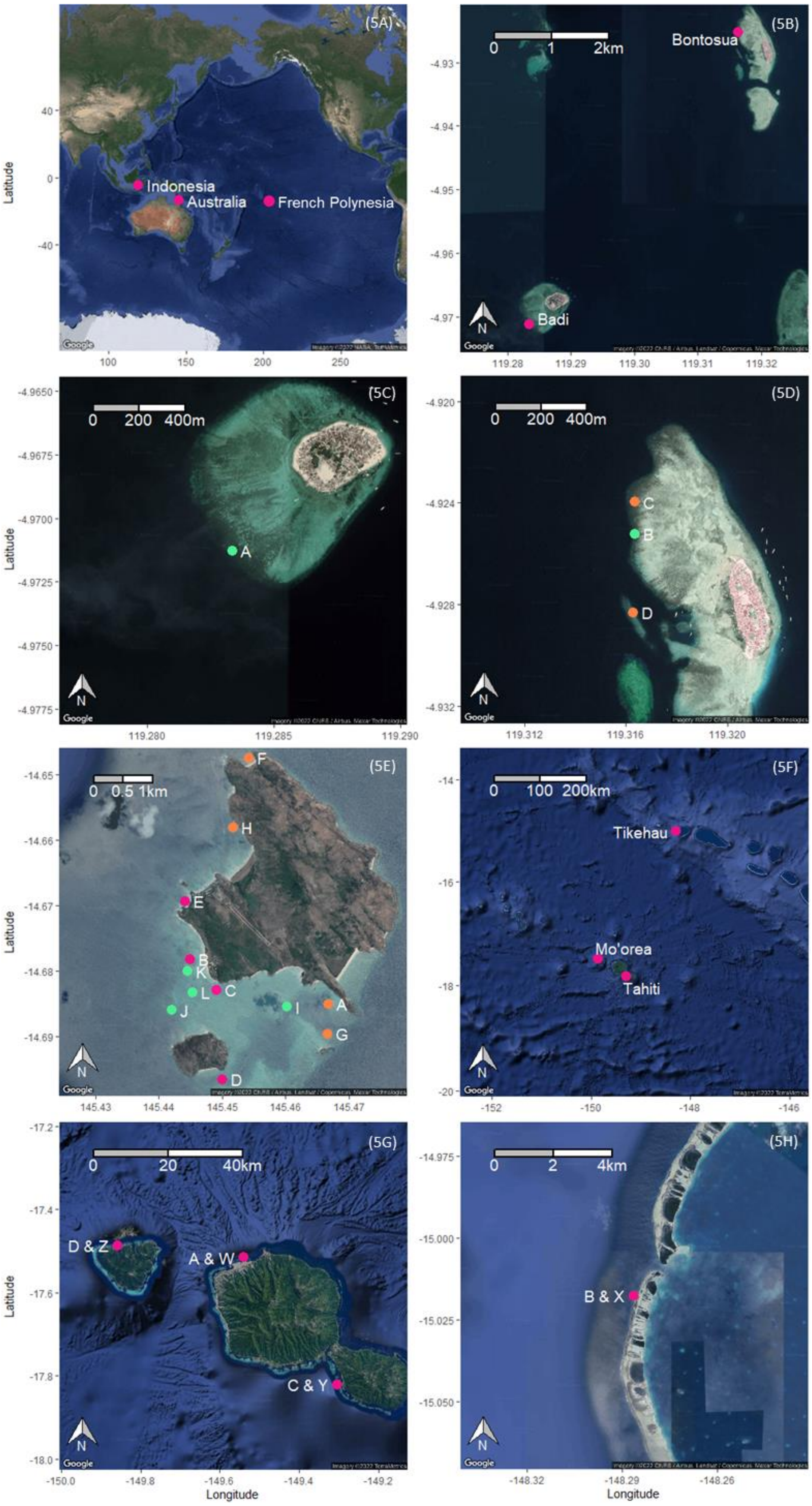


**Fig. S5.** Map of study locations and sites. (A) The location where each dataset was collected. **(B)** The location of Bontosua and Badi islands, where study sites in the Indonesian dataset were located. **(C)** The location of the study site on Badi Island. **(D)** The location of the sites around Bontosua Island. Healthy and degraded sites around Bontosua and Badi Islands are labelled in green and orange respectively. **(E)** The location of study sites around Lizard Island, where the Australian dataset was collected. High fish diversity sites are labelled in green, low fish diversity sites are labelled in orange and four sites excluded from ecological category tasks are in pink. **(F)** The location of Mo'orea, Tahiti and Tikehau, where study sites in the French Polynesian dataset were located. **(G)** The location of study sites around Mo'orea and Tahiti. **(H)** The location of the study site on Tikehau.

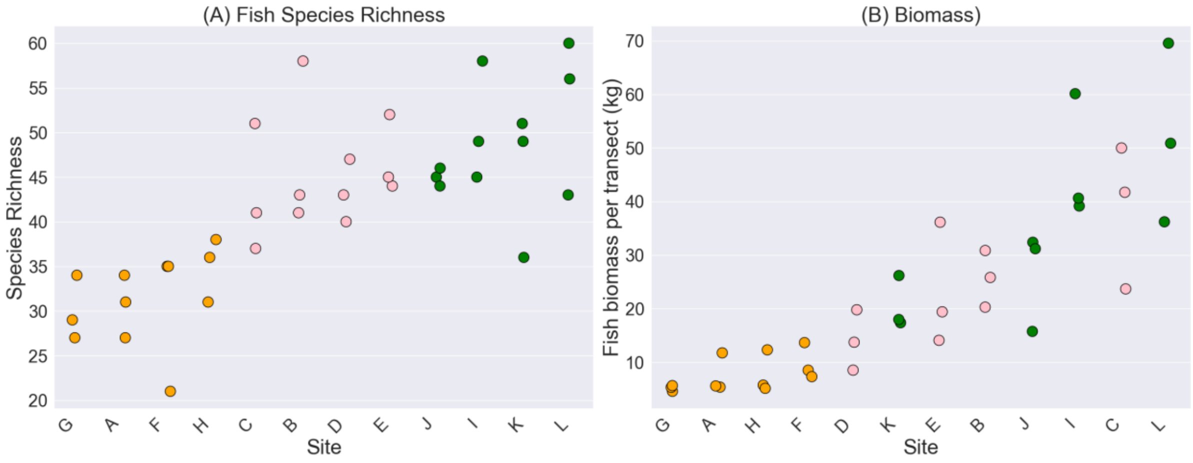


**Fig. S6.** Plots showing (A) fish species richness and (B) total fish assemblage biomass recorded from three transect surveys on each of the 12 sites around Lizard Island, Australia. Two ecological categories were first created using the four highest and four lowest scoring sites for species richness, marked in green and orange respectively. These two categories were also found to have non-overlapping biomass values and therefore the categories were labelled as ‘high fish diversity’ and ‘low fish diversity’ sites. The four sites excluded from ecological category tasks are labelled in pink.

### Supplementary tables

Including: Table S1 raw classifier results; Table S2 ANOVA results from comparing classifier accuracies; Table S3 details on acoustic indices used to construct the compound index.

**Table S1.** Mean and standard deviation of classifier accuracy across repeated training instances using each the three machine learning methods (compound index, pretrained CNN and trained CNN) at six different tasks. Accuracy is the proportion of one-minute recordings from the test data that were correctly classified. Accuracy with random assigment indicates the expected accuracy of a model that performs random classification. N = 100 for all tasks, except the 'Fish rich or poor (Australia)' and 'Shallow or mesophotic (French Polynesia)' tasks, where N = 32. Methods where accuracy was reported as significantly higher by the ANOVA test are indicated in superscript next to the mean value for the respective method. Here, the letter denotes which of the other two methods the selected method was higher than ('A' = compound index, 'B' = pretrained CNN and 'C' = CNN).

| | Compound index | | Pretrained CNN | | Trained CNN | | Accuracy using random assignment |
|---|---|---|---|---|---|---|---|
| Task | Mean | Standard deviation | Mean | Standard deviation | Mean | Standard deviation | |
| High or low coral cover (Indonesia) | 0.86 | 0.10 | 0.88 | 0.08 | $0.91^{AB}$ | 0.09 | 0.50 |
| Site identification (Indonesia) | 0.85 | 0.09 | 0.85 | 0.08 | $0.89^{AB}$ | 0.08 | 0.25 |
| High or low fish diversity (Australia) | 0.71 | 0.12 | 0.72 | 0.09 | 0.73 | 0.11 | 0.50 |
| Site identification (Australia) | $0.56^{B}$ | 0.05 | 0.52 | 0.07 | 0.54 | 0.09 | 0.08 |
| Shallow or mesophotic (French Polynesia) | 0.88 | 0.16 | 0.82 | 0.20 | 0.83 | 0.20 | 0.25 |
| Site identification (French Polynesia) | $0.99^{B}$ | 0.00 | 0.99 | 0.00 | $1.00^{AB}$ | 0.00 | 0.13 |

**Table S2**. Three-way ANOVAs comparing supervised classifier accuracy over six different tasks across repeated training instances for the three machine learning methods (compound index, pretrained CNN and trained CNN). Where significant differences ($p<0.05$) were detected, post hoc Tukey HSD tests were performed, otherwise cells are left empty. Under the Tukey HSD heading, entries under the first sub-column indicate this method reported a significantly higher accuracy than the method in the second sub-column beneath. 95% Confidence intervals (CI) represent the range of estimated differences in classifier accuracy between the respective pair of methods.

| | | **ANOVA result** | | **Tukey HSD results** | | | | | |
|---|---|---|---|---|---|---|---|---|---|
| | | | | **Compound index** | | **Pretrained CNN** | | **CNN** | |
| **Dataset** | **Task** | **f-value** | **p-value** | **Pretrained CNN** | **CNN** | **Compound index** | **CNN** | **Compound index** | **Pretrained CNN** |
| Indonesia | Habitat: high or low coral cover | 9.825 | <0.001 | | | | | p = 0.001, CI = 0.026 to 0.085 | p = 0.027, CI = 0.003 to 0.062 |
| Indonesia | Site identification | 8.939 | <0.001 | | | | | p = 0.003, CI = 0.012 to 0.067 | p = 0.001, CI = 0.019 to 0.074 |
| Australia | Habitat: fish rich or poor | 0.180 | 0.836 | | | | | | |
| Australia | Site identification | 7.495 | 0.001 | p = 0.001, CI = 0.015 to 0.063 | | | | | |
| French Polynesia | Habitat: shallow or mesophotic | 1.143 | 0.323 | | | | | | |
| French Polynesia | Site identification | 268.343 | <0.001 | p = 0.011, CI = 0.0003 to 0.002 | | | | p = 0.001, CI = 0.007 to 0.0095 | p = 0.011, CI = 0.009 to 0.01 |

**Table S3.** Acoustic indices selected. All indices were calculated using the scikit-maad package (v1.3). Where additional settings are 'None', the index was calculated over the precomputed spectrogram with no further parameters required. For spectrogram calculation a hanning window, window length (nperseg) of 256, and, power spectral density mode were set using spectrogram function in the packages 'sound' module. The reference column cites a study which has reported a relationship between the respective index and some aspect of coral reef ecology.

| Index | Mechanism | Additional settings | Reference |
|---|---|---|---|
| Acoustic Complexity Index (ACI) | Measures variability in intensity of frequencies across time | None | Bertucci et al., Sci Rep (2016). |
| Acoustic Diversity Index (ADI) | Measures diversity across frequency bands | Min and max frequencies matched the frequency band in use. bin_step was 1/10th of the bands range. dB threshold = -50. | Williams et al,. Eco Ind (2022). |
| Acoustic Entropy (H) | Measures randomness across temporal and spectral domains | s = QUT, mode = fast, Nt = 256 | Bertucci et al., Sci Rep (2016). |
| Amplitude Index (M) | Measures median of amplitude envelope | mode = fast, Nt = 256 | Williams et al,. Eco Ind (2022). |
| Bioacoustic Index (BI) | Measures cumulative intensity across frequency bands | Min and max frequencies matched the frequency band in use | Elise et al,. Eco Ind (2019). |
| Normalised mean difference index (NDSI) | Measures amplitude difference between two selected frequency bands | flim_bioph = (2000, 8000), flim_antroPh = (50, 2000) | Elise et al,. Rem Sen (2022). |
| Spectral entropy (Hf) | Measures randomness across the frequency domain | None | Elise et al,. Eco Ind (2019). |
| Temporal Entropy (Ht) | Measures randomness across the temporal domain | mode = fast, Nt = 256 | Elise et al,. Eco Ind (2019). |
